# The E3 ligase TRIM1 ubiquitinates LRRK2 and controls its localization, degradation, and toxicity

**DOI:** 10.1101/2020.10.21.336578

**Authors:** Adrienne E. D. Stormo, Farbod Shavarebi, Molly FitzGibbon, Elizabeth M. Earley, Hannah Ahrendt, Lotus S. Lum, Erik Verschueren, Danielle L. Swaney, Gaia Skibinski, Abinaya Ravisankar, Jeffrey van Haren, Emily J. Davis, Jeffrey R. Johnson, John Von Dollen, Carson Balen, Jacob Porath, Claudia Crosio, Christian Mirescu, Ciro Iaccarino, William T. Dauer, R. Jeremy Nichols, Torsten Wittmann, Timothy C. Cox, Steve Finkbeiner, Nevan J. Krogan, Scott A. Oakes, Annie Hiniker

## Abstract

Missense mutations in leucine-rich repeat kinase 2 (LRRK2) are the most common cause of familial Parkinson’s Disease (PD); however, pathways regulating LRRK2 subcellular localization, function, and turnover are not fully defined. We performed quantitative mass spectrometry-based interactome studies to identify 48 novel LRRK2 interactors, including the microtubule-associated E3 ubiquitin ligase TRIM1 (<u>Tri</u>partite <u>M</u>otif Family 1). TRIM1 recruits LRRK2 to the microtubule cytoskeleton for ubiquitination and proteasomal degradation by binding LRRK2_911-920_, a nine amino acid segment within a flexible interdomain region (LRRK2_853-981_), which we designate the “Regulatory Loop” (RL). Phosphorylation of LRRK2 Ser910/Ser935 within LRRK2 RL serves as a molecular switch controlling LRRK2’s association with cytoplasmic 14-3-3 versus microtubule-bound TRIM1. Association with TRIM1 modulates LRRK2’s interaction with Rab29 and prevents upregulation of LRRK2 kinase activity by Rab29 in an E3-ligase-dependent manner. Finally, TRIM1 rescues neurite outgrowth deficits caused by PD-driving mutant LRRK2 G2019S. Our data suggest that TRIM1 is a critical regulator of LRRK2, controlling its degradation, localization, binding partners, kinase activity, and cytotoxicity.

## Introduction

Leucine-rich repeat kinase 2 (LRRK2) mutations are the most common genetic cause of Parkinson’s Disease (PD), a devastating neurodegenerative disorder affecting 1-2% of people over age 65.^1, 2^ LRRK2 is a 290 kDa polypeptide with multiple protein-protein interaction domains—including N-terminal armadillo, ankyrin, and LRR domains, and C-terminal WD40 repeats—that flank enzymatically active Roc GTPase (Ras of complex proteins), COR, and kinase domains (Figure 1a). Several point mutations in the catalytic core of LRRK2 cause autosomal dominant PD with incomplete penetrance (referred to herein as “LRRK2-PD”), while other mutations in the protein increase risk for sporadic PD.^3, 4^ The most common LRRK2-PD mutation, LRRK2 G2019S, falls in the kinase domain, as does the I2020T mutation. Several other PD-driving mutations, including R1441G/C/H and Y1699C, are located in the Roc and COR domains and promote GTP binding.^5, 6^ A distinct set of LRRK2 mutations augments risk for Crohn’s disease, leprosy, and tuberculosis.^7–10^ How LRRK2 mutations cause PD is not well understood; however, mounting evidence supports a toxic gain-of-function mechanism with PD-driving LRRK2 mutations demonstrating abnormally augmented kinase activity.^11, 12^ Certain PD mutants have also been shown to change LRRK2’s affinity for binding partners (such as R1441 mutants which do not bind 14-3-3 proteins) and others have been suggested to increase LRRK2 protein levels.^11, 13, 14^ LRRK2 kinase inhibition is being pursued as a possible therapeutic avenue for PD, with highly selective kinase inhibitors in clinical trials for LRRK2-driven and sporadic PD. An alternative approach that has not been explored is to reduce LRRK2 activity by exploiting cellular pathways that augment LRRK2 degradation, thus decreasing total LRRK2 protein levels.

**Figure 1.**
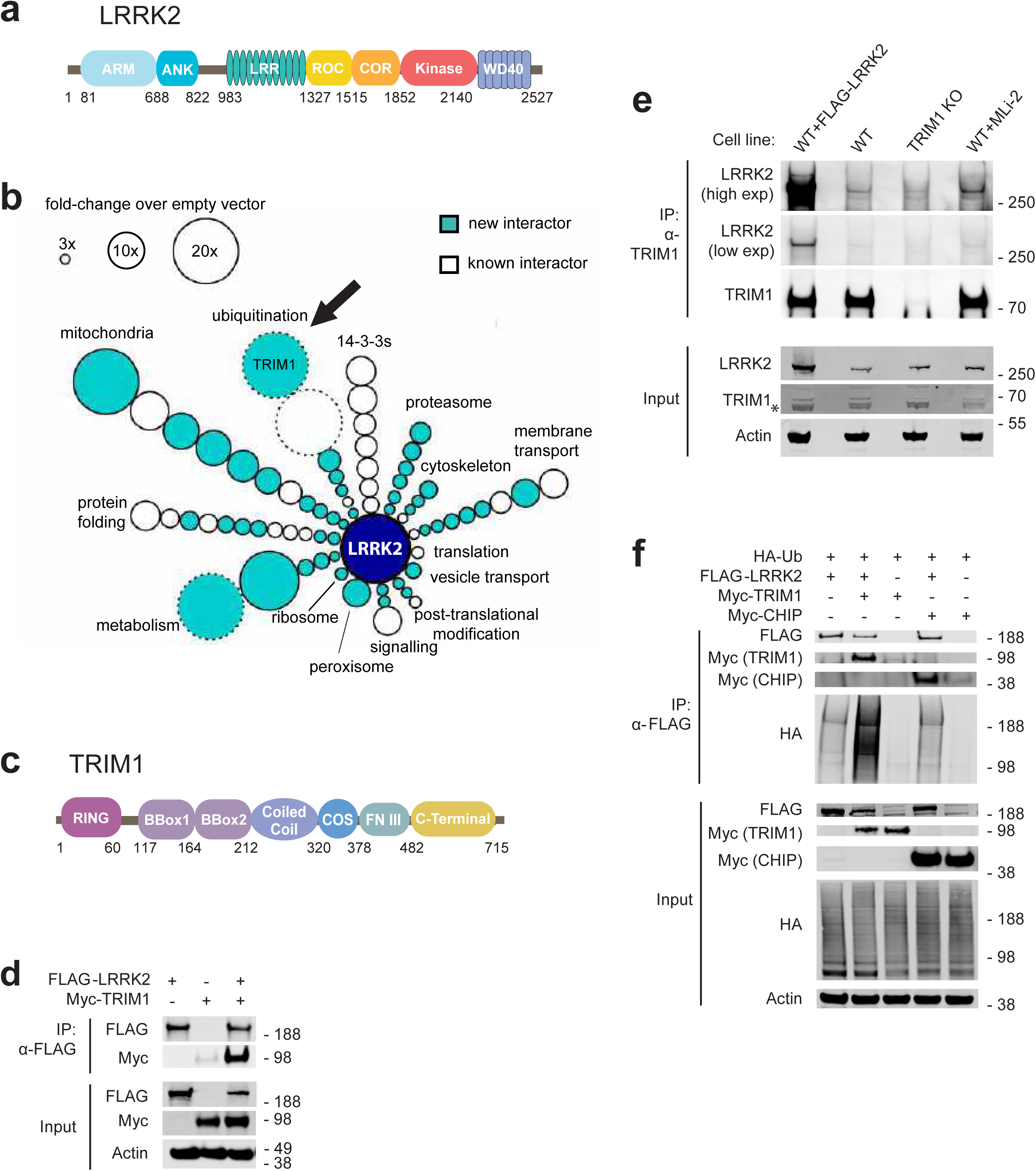
TRIM1 is a new LRRK2 E3 ubiquitin ligase. (A) Diagram of LRRK2 protein domains (ARM: armadillo repeat; ANK: ankyrin repeat; LRR: leucine-rich repeat; ROC: ras of complex proteins; COR: C-terminal of ROC domain). (B) Schematic of LRRK2 interactome in HEK-293T cells. LRRK2 interacting partners are classified radially according to function (aqua: new LRRK2 interacting partners, white: previously identified LRRK2 partners, size of circle indicates fold-change over empty vector control, circles without solid black outline had no peptides present in empty vector control, arrow indicates TRIM1). FLAG-LRRK2 was immunoprecipitated and interacting partners identified and quantified by MS. Data represent at least four total independent replicates from two experiments and are additionally shown in table S1. (C) Diagram of TRIM1 protein domains (FNIII: fibronectin III domain). (D) Co-immunoprecipitation of myc-TRIM1 with FLAG-LRRK2 in HEK-293T cells. (E) Co-immunoprecipitation of endogenous LRRK2 with TRIM1 in WT HEK-293T and HEK-293T TRIM1 CRISPR knock-out line. From left to right: WT HEK-293T cells transfected with exogenous FLAG-LRRK2 (positive control), WT HEK-293T cells, TRIM1 knockout HEK-293T cells, WT HEK-293T cells treated with 500 nM MLi-2 for five hours. “Low exp” indicates short exposure of membrane, “high exp” indicates longer exposure of membrane. (F) Co-immunoprecipitation and ubiquitination of FLAG-LRRK2 with myc-TRIM1 or myc-CHIP in the presence of HA-ubiquitin in HEK-293T cells.

The specific pathways regulating LRRK2 protein degradation are only beginning to be comprehensively evaluated, and both the autophagic-lysosome and ubiquitin-proteasome systems appear to be involved. LRRK2 has a complex relationship with autophagy: multiple studies implicate LRRK2 in regulating autophagy,^15, 16^ and a portion of LRRK2 appears to be degraded by chaperone-mediated autophagy.^17^ Additionally, a large fraction of endogenous LRRK2 has been shown to be degraded via the ubiquitin-proteasome system;^18^ however, thus far, only two proteins are reported to act as E3 ubiquitin ligases for LRRK2: (1) WD repeat and SOCS box containing 1 (WSB1), which ubiquitinates LRRK2 via atypical K27 and K29 linkages and causes LRRK2 aggregation but does not appear to promote proteasomal degradation;^19^ and (2) C-terminus of Hsc70-interacting protein (CHIP), an HSP70 co-chaperone that interacts with many partially folded proteins.^20, 21^ In keeping with its preference for misfolded proteins,^22^ CHIP appears to be particularly important for turnover of destabilized LRRK2 variants, such as the sporadic PD modest risk allele LRRK2 G2385R, and may not be as critical for other LRRK2-PD mutants.^23^

LRRK2 is present at low levels in most cells types, hindering definitive determination of its endogenous subcellular localization. Predominantly through the use of overexpression systems, LRRK2 has been found (1) associated with endolysosomal and golgi membranes, where it interacts with Rab GTPases;^24^ (2) present in the cytoplasm, where it binds the 14-3-3 family of adapter proteins;^25^ and (3) at the cytoskeleton, where it interacts with microtubules.^26, 27^ Strong evidence demonstrates that LRRK2 associates with membranes; important recent work identified 14 membrane-associated Rab proteins as kinase substrates of LRRK2, including Rab10 and Rab29.^28^ Rab29, which localizes to Golgi network membranes, was also shown to be a unique activator of LRRK2’s kinase activity, at least in cellular overexpression systems, as measured by LRRK2 autophosphorylation at Ser1292 and phosphorylation of substrate Rabs.^24, 29^ Rab29 appears to preferentially activate Roc-COR domain LRRK2-PD mutants such as LRRK2 R1441G.^24^ The armadillo domain as well as conserved Leu-rich motifs in the ankyrin domain of LRRK2 have been shown to be important for LRRK2 to bind Rab29.^24, 30, 31^

LRRK2 also localizes to the cytoplasm, where it associates with 14-3-3 proteins, a family of seven highly homologous isoforms that function as adaptor proteins to regulate myriad signaling pathways.^32^ The structural features mediating LRRK2’s interaction with 14-3-3 have been rigorously investigated and phosphorylation of LRRK2 serine residues Ser910 and Ser935 is required.^33^ LRRK2-PD mutants with decreased phosphorylation of Ser910 and Ser935 (predominantly species with mutations in the Roc-COR domain) show reduced affinity for 14-3-3 proteins.^13, 33, 34^ LRRK2’s interaction with 14-3-3 appears necessary to maintain LRRK2 in the cytoplasm and may be one mechanism that prevents Rab29-mediated LRRK2 kinase activation. In support of this model, abnormal LRRK2 function was recently implicated in idiopathic PD (i.e., PD negative for LRRK2 mutations): sensitive proximity ligation assays were used to demonstrate both increased kinase activity and decreased 14-3-3 binding of LRRK2 in substantia nigra neurons from patients with idiopathic PD compared to controls, strengthening the link between abnormalities in LRRK2 function and idiopathic PD.^35^

A growing body of evidence indicates that LRRK2 can also associate with the microtubule cytoskeleton. Multiple groups have demonstrated that LRRK2 forms filamentous structures along microtubules,^26, 27, 36^ which increase in frequency with kinase inhibitor treatment or point mutations in either the Roc-COR or kinase domains.^26, 27, 36^ Very recently, the *in situ* cryo-electron tomography structure of a truncated variant of PD-mutant LRRK2 I2020T bound to microtubules was solved to 14 Å, showing the Roc-COR domain adjacent to the microtubule and the kinase domain exposed to the cytoplasm.^37^ In keeping with this structure, LRRK2 can directly interact with β-tubulin through its Roc domain, altering tubulin acetylation and inhibiting axonal transport in neurons.^38–40^ A second recent work, which solved the structure of LRRK2 to 3.5 Å using cryo-EM, suggests that LRRK2’s interaction with microtubules is regulated by the conformation of its kinase domain and further that LRRK2 binding to microtubules can disrupt axonal transport.^41^ Axonal transport is restored by increasing microtubule acetylation, suggesting that the LRRK2-microtubule interaction is regulated and occurs only at specific subtypes of microtubules.^39^ However, additional upstream signals or binding partners regulating LRRK2 localization to microtubules have not previously been identified.

Here, we used a mass spectrometry (MS) interactome approach to find new LRRK2 binding partners and discovered the little-studied, microtubule-localized E3 ubiquitin ligase TRIM1. While 14-3-3 stabilizes LRRK2 in the cytoplasm and Rab29 augments LRRK2’s kinase activity at membranes, TRIM1 recruits LRRK2 to the microtubule cytoskeleton, where it mediates LRRK2 ubiquitination and proteasomal degradation. We narrowed down the TRIM1 binding site to nine amino acids (911-920) within a flexible interdomain (“Regulatory Loop” or RL) region of LRRK2, LRRK2_853-981_. LRRK2 RL contains Ser 910 and Ser 935, and the phosphorylation status of these serine residues influences LRRK2’s choice of binding partner (14-3-3 versus TRIM1). Finally, TRIM1 inhibits Rab29-mediated activation of LRRK2 kinase activity and rescues PD-mutant LRRK2-driven toxicity as measured by neurite outgrowth. Our studies show that TRIM1 is an important E3 ligase influencing LRRK2 subcellular location, protein levels, and function. They also suggest that LRRK2’s RL is a critical structural element whose post-translational modification is important in controlling LRRK2’s binding to interacting partners, which in turn regulates LRRK2 localization, turnover, kinase function, and toxicity.

## Results

### TRIM1 is a novel LRRK2 E3 ubiquitin ligase

We postulated that critical LRRK2 partners may have been missed in previous interaction studies, some of which used only a portion of the protein as bait.^38, 40, 42, 43^ We used an established affinity purification-MS (AP-MS) approach, which has been extensively validated in our lab, to systematically and quantitatively identify the interactome of full-length LRRK2.^44^ N-terminally FLAG-tagged full-length LRRK2 or FLAG-alone control plasmids were transiently transfected into HEK-293T cells (selected for our extensive library of baseline interactome data, allowing better exclusion of non-specific interactors); lysates were affinity purified; and the eluted material subjected to MS as in Jager *et al.*^44^ Interacting partners were determined by label-free MS1 quantification using MSStats.^45^ High-confidence interaction partners were proteins with an intensity >3-fold increased over empty vector control (p-value <0.05), which identified >20 previously reported LRRK2-interacting proteins, including all members of the 14-3-3 family of proteins, as well as 48 novel partners, which were categorized according to function (Figure 1b and Table S1). The top hit was the putative E3 ubiquitin ligase, tripartite motif family 1 (TRIM1, also called MID2), which has never been described as playing a role in LRRK2 biology, though a prior proteomics study did identify TRIM1 as a possible LRRK2 interacting partner in HEK-293T cells.^42^

TRIM1 is a little-studied 78 kDa protein whose coding sequence is located on the X-chromosome within the PARK12 genomic locus.^46^ It is a member of a ∼75 protein superfamily of E3 ubiquitin ligases with a common tripartite motif consisting of a RING domain, one or two B-box-type zinc fingers, and a coiled-coil domain.^47^ TRIM1’s tripartite motif is followed by a microtubule-targeting COS domain, a fibronectin type III domain, and a C-terminal domain (Figure 1c).^48^ While its cellular functions remain largely uncharacterized, TRIM1 missense mutations were reported in families with a rare form of X-linked mental retardation, indicating a critical role in normal brain function.^49^ Consistent with our MS findings, myc-TRIM1 and FLAG-LRRK2 overexpressed in HEK-293T cells robustly co-immunoprecipitated (Figure 1d).

To validate the endogenous interaction of LRRK2 with TRIM1, we generated a TRIM1 knockout (KO) HEK-293T cell line using CRISPR/Cas9 gene editing. Genomic sequencing of the TRIM1 KO line identified two N-terminal frame shift mutations in the TRIM1 gene leading to stop codons at amino acids 14 and 18 and no wild type alleles of TRIM1. Because TRIM1 is expressed at low levels endogenously, it is not visible on immunoblot of HEK-293T cell lysates (Figure 1e, asterisk (*) in TRIM1 immunoblot of lysate indicates a non-specific band). However, endogenous TRIM1 is detectable upon immunoprecipitation from HEK-293T cells. Immunoprecipitation of endogenous TRIM1 protein showed absence of TRIM1 protein in the TRIM1 KO line in contrast to the WT line. Overexpressed FLAG-LRRK2 robustly co-immunoprecipitated with endogenous TRIM1 (Figure 1e, lane 1). Endogenous LRRK2 co-immunoprecipitated with endogenous TRIM1 in the TRIM1 WT line, in contrast to the TRIM1 KO line (Figure 1e, compare lanes 2 and 3). We noted that this interaction was enhanced by addition of the LRRK2 kinase inhibitor MLi-2 for five hours prior to immunoprecipitation (Figure 1e, compare lanes 2 and 4). Our data demonstrate that TRIM1 and LRRK2 interact under both overexpression and endogenous conditions.

Given that many TRIM family members are RING-finger E3 ubiquitin ligases, we speculated that TRIM1 may function to ubiquitinate LRRK2. We utilized an established *in vivo* ubiquitination assay for LRRK2, which was previously used to demonstrate LRRK2 ubiquitination by CHIP (the single E3 ubiquitin ligase known to target LRRK2 for proteasomal degradation).^20^ We found that coexpression of myc-TRIM1 with FLAG-LRRK2 and HA-ubiquitin resulted in robust LRRK2 ubiquitination with myc-CHIP serving as a positive control and HA-ubiquitin alone serving as a negative control (Figure 1f). Thus, TRIM1 is a novel E3 ubiquitin ligase for LRRK2.

### TRIM1 recruits LRRK2 to microtubules

TRIM1 is part of the six-member C-I subfamily of TRIM proteins, all of which strongly associate with microtubules through a C-terminal COS domain.^48^ Therefore, we hypothesized that LRRK2 may interact with TRIM1 at the microtubule cytoskeleton. Using overexpression studies, multiple groups have identified a portion of LRRK2 at microtubules;^26, 38^ however, the fraction of LRRK2 associated with microtubules is generally very low in the absence of LRRK2 point mutations or kinase inhibitors.^27, 39^ We used live cell confocal microscopy to examine the subcellular distribution of transfected full-length GFP-LRRK2 in human lung H1299 cells, chosen for their large size and flat shape, allowing clear evaluation of the microtubule network. In agreement with previous studies, the vast majority of GFP-LRRK2 was cytoplasmic and did not co-localize with microtubules labeled with mCherry-tubulin (Figure 2a). We confirmed previous work demonstrating that mCherry-TRIM1 localizes to microtubules (Figure 2b).^48, 50, 51^ Strikingly, co-expression of mCherry-TRIM1 substantially increased GFP-LRRK2 co-localization with microtubules (Figure 2b and Movie S1). We observed that mCherry-TRIM1 recruited GFP-LRRK2 to microtubules in all cell lines examined, including human lung carcinoma (A549) cells (Figure S1a), human neuroblastoma (SK-N-SH) cells (Figure S1b), and human embryonic kidney (HEK-293T) cells (Figure S1c), as well as human breast carcinoma (MCF7), and human (SH-SY5Y) and mouse (N2a) neuroblastoma cells (not shown). Microtubule-localized GFP-LRRK2 showed a discontinuous appearance, in keeping with observations from other groups.^27^ Quantification of the percentage of cells with microtubule-associated GFP-LRRK2 in the presence of mCherry-TRIM1 versus mCherry-tubulin (Figure 2F) revealed that TRIM1 caused LRRK2 microtubule localization in essentially all cells in which both proteins were expressed (98.7% +/- 1.1%, mean +/- standard deviation) while only rare cells expressing mCherry-tubulin had visible LRRK2 at microtubules (6.3% +/- 2.1%, mean +/- S.D.).

**Figure 2.**
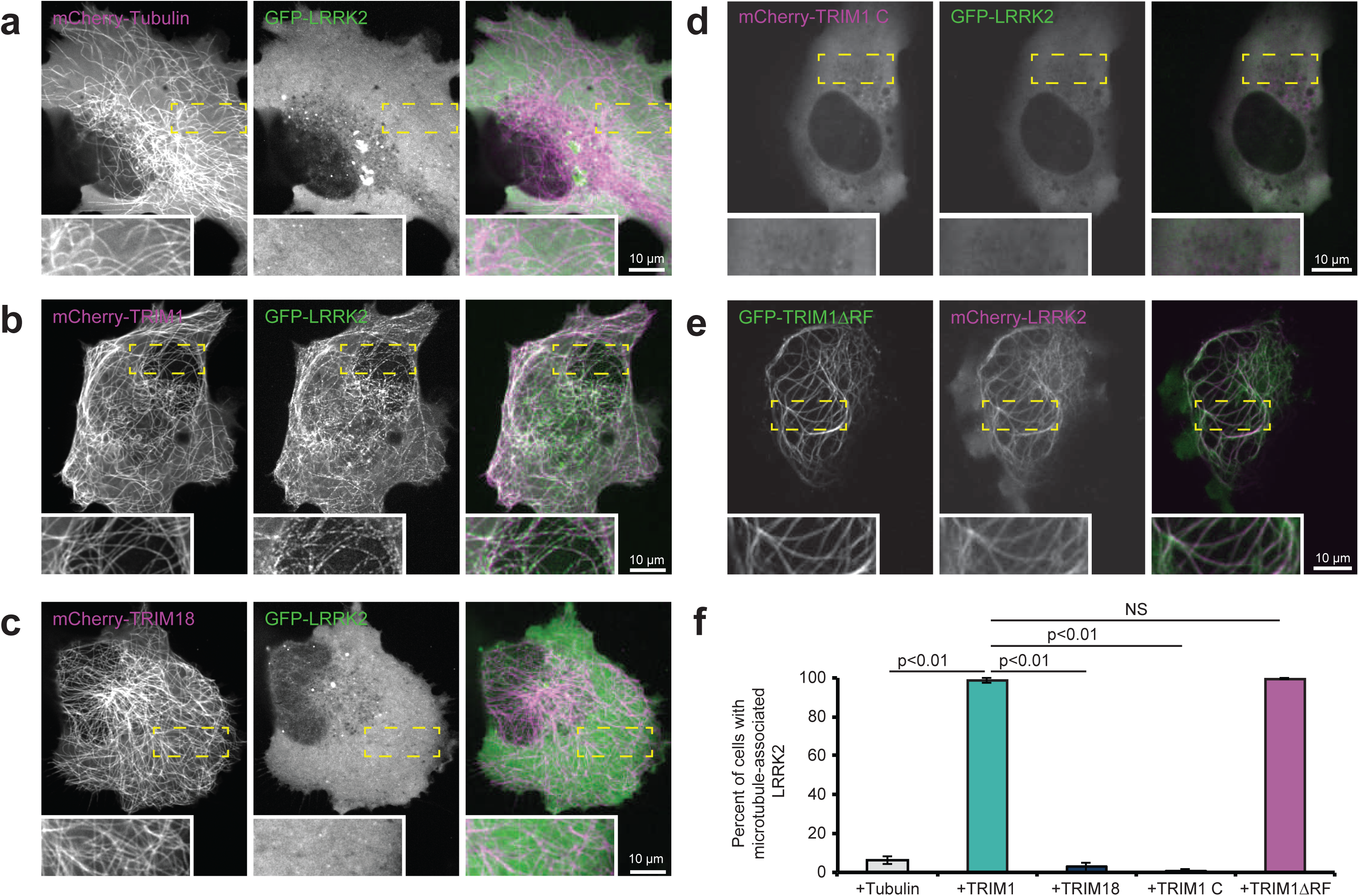
TRIM1 co-expression recruits LRRK2 to microtubules. Live-cell confocal microscopy of fluorescently tagged LRRK2 and tubulin, TRIM1, or TRIM18 constructs transiently transfected into H1299 cells. Insets in A-E show higher magnification of region identified by the yellow box. (A) In the presence of mCherry-tubulin, GFP-LRRK2 is diffusely cytoplasmic. From left to right: mCherry-tubulin, GFP-LRRK2, merged image. (B) In the presence of mCherry-TRIM1, GFP-LRRK2 localizes to microtubules. From left to right: mCherry-TRIM1, GFP-LRRK2, merged image. (C) In the presence of mCherry-TRIM18, GFP-LRRK2 is diffusely cytoplasmic. From left to right: mCherry-TRIM18, GFP-LRRK2, merged image. (D) mCherry-TRIM1 C is cytoplasmic. When co-expressed with GFP-LRRK2, both remain diffusely cytoplasmic. From left to right: mCherry-TRIM1 C, GFP-LRRK2, merged image. (E) GFP-TRIM1 ΔRF maintains microtubule localization and co-localizes with mCherry-LRRK2. From left to right: GFP-TRIM1 ΔRF, mCherry-LRRK2, merged image. (F) Quantification of cells with microtubule-associated LRRK2 when co-expressed with indicated proteins in H1299 cells. 100 cells were evaluated in each condition in each of three independent experiments, bars show mean +/- standard deviation. Significance testing for panel F was performed using Kruskal-Wallis with post-hoc Dunn test and Bonferroni correction. Scale bars = 10 µm.

We next evaluated the specificity of the TRIM1-LRRK2 interaction in controlling LRRK2 microtubule localization. Of the ∼75 members of the TRIM family, TRIM1 is most homologous to TRIM18 (76% identical, 88% similar, Figure S1d). Like TRIM1, TRIM18 binds microtubules. Loss-of-function mutations in TRIM18 cause a syndrome of congenital midline defects (X-linked Opitz G/BBB syndrome), which has not been observed for TRIM1 mutations.^50^ Co-expression of mCherry-TRIM18 was insufficient to recruit LRRK2 to microtubules (Figure 2c). The percentage of cells with GFP-LRRK2 at microtubules in the presence of mCherry-TRIM18 was 2.8% +/- 2.5%, which is statistically indistinguishable from that of mCherry-tubulin. Additionally, myc-TRIM18 did not robustly co-immunoprecipitate with FLAG-LRRK2 and its ability to ubiquitinate FLAG-LRRK2 was much diminished compared to myc-TRIM1 (Figure 3a). We also tested the ability of GFP-LRRK2 to bind TRIM9, which has the same domain organization as TRIM1 and TRIM18 (25% identical, 39% similar) and has been linked to PD in one study.^52^ As with TRIM18, myc-TRIM9 failed to appreciably co-immunoprecipitate with LRRK2 (Figure S1e). Thus, the LRRK2-TRIM1 interaction appears highly specific.

**Figure 3.**
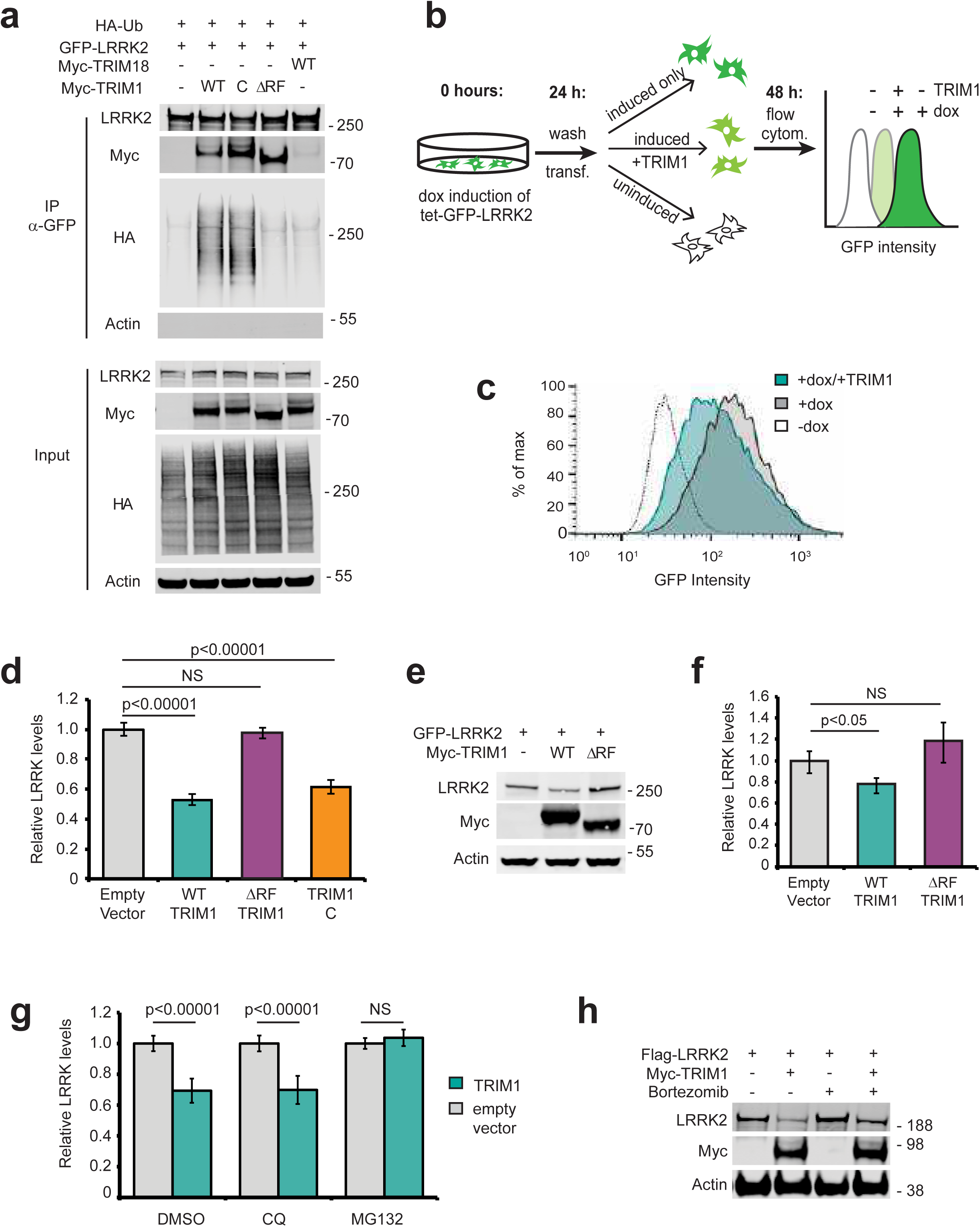
TRIM1 ubiquitinates LRRK2 to regulate its proteasomal degradation. (A) Immunoprecipitation and ubiquitination of GFP-LRRK2 with myc-TRIM1, myc-TRIM1 C, myc-TRIM1 ΔRF, or myc-TRIM18 in the presence of HA-ubiquitin in HEK-293T cells. (B) Schematic of flow cytometric assay using GFP fluorescence to measure GFP-LRRK2 turnover. Doxycycline-inducible GFP-LRRK2 HEK-293T cells are induced for 18 to 24 hours, transfected and doxycycline simultaneously withdrawn, and GFP fluorescence measured after 18-24 hours (additional validation of assay in Figure S2). All flow cytometry assays were performed in the doxycycline-inducible GFP-LRRK2 HEK-293T cell lines described in Zhao *et al.*^62^ (C) Representative histograms of GFP-LRRK2 fluorescence in the absence or presence of doxycycline followed by TRIM1 or empty vector transfection. (D) Quantification of GFP-LRRK2 levels 24 hours after doxycycline withdrawal in the presence of empty vector (grey bar), TRIM1 (green bar), TRIM1 ΔRF (purple bar), or TRIM1 C (orange bar). (E) Representative immunoblot of GFP-LRRK2 levels from dox-inducible HEK-293T cells in the presence of myc-TRIM1 WT, myc-TRIM1 ΔRF, or empty vector. (F) Quantification of (E) showing mean value with error bars showing the standard error of the mean. (G) Quantification of GFP-LRRK2 levels in the presence of chloroquine (CQ) at 25 μM for 24 hours, MG132 at 2 μM for 24 hours, or equivalent volume of DMSO vehicle. (H) Immunoblot of FLAG-LRRK2 levels with or without expression of myc-TRIM1 in the absence or presence of proteasomal inhibitor bortezomib (1 nM for 18 hours). Bar graphs of flow cytometry assays (panels D and G) show normalized median green fluorescence intensity with error bars showing twice the standard error of the mean. All histograms and bar charts of flow cytometry results represent at least 10,000 single cells per condition. All co-immunoprecipitation and flow cytometry assay results show a representative experiment, with the experiment repeated a minimum of 3 times. Significance for flow cytometry data (panels D and G) are calculated using ANOVA with post-hoc t-test with Bonferroni correction. Significance testing for panel F was performed using Mann-Whitney U test.

To evaluate the extent to which TRIM1’s microtubule-binding function is required for its E3 ligase activity, we constructed two variants of TRIM1, one which does not localize to microtubules and is instead cytoplasmic, and the other which lacks the RING domain, eliminating its E3 ligase function. To construct the cytoplasmic variant of TRIM1, we utilized previous work on TRIM18 showing that mutating six amino acids in TRIM18’s COS domain to alanine prevents TRIM18 from binding to microtubules and redirects it to the cytoplasm.^48^ The identical amino acids are present in TRIM1 and were mutated to alanine (FLQ328AAA LDY377AAA, Figure S1d). The resulting construct, which we call TRIM1 C (for cytoplasmic), is diffusely cytoplasmic (Figure 2d) but retains E3 ligase activity and LRRK2 binding (Figure 3a). The RING finger deleted TRIM1 (“TRIM1 ΔRF”) is microtubule-bound (Figure 2e) but does not show E3 ligase activity (Figure 3a). The percentage of cells with GFP-LRRK2 at microtubules in the presence of mCherry-TRIM1 C was 0.7% +/- 1.1%. The percentage of cells with mCherry-LRRK2 at microtubules in the presence of GFP-TRIM1 ΔRF was 99.7 % +/- 0.6%. Therefore, TRIM1’s ability to ubiquitinate LRRK2 can be separated from its ability to localize LRRK2 to the microtubule network.

Repeated attempts under multiple experimental conditions did not allow us to visualize the subcellular localization of endogenous LRRK2. Live cell imaging was performed using A549 cells with an N-terminal GFP-tag added to *LRRK2* by CRISPR editing (gift of Dario Alessi, unpublished). Immunofluorescence using a variety of commercially available antibodies against LRRK2 (MJFF C41-2, UDD3, N231) and GFP (Abcam 13970) was performed on the GFP-LRRK2 A549 line as well as wild type versus CRISPR *LRRK2* knockout A549 cells (gift of Dario Alessi), wild type versus TALEN *LRRK2* knockout murine RAW 264.7 macrophages (from the Michael J. Fox Foundation), and wild type compared to siRNA *LRRK2* knockdown human melanoma Malme-3M cells. Under no experimental condition could we visualize a fluorescence signal specific to endogenous LRRK2. The inability to reproducibly visualize endogenous LRRK2 using these methods is in keeping with previous reports and highlights an important limitation in the field.^53^

### TRIM1 ubiquitinates LRRK2 to regulate its turnover via the proteasome

Polyubiquitin linkages frequently serve to target proteins for proteasomal or autophagic degradation, though they may also signal other molecular functions.^54^ Co-expression of TRIM1 with LRRK2 decreased LRRK2 accumulation over time compared to co-expression of control vector (Figure S2a, quantified in S2b), suggesting TRIM1-mediated ubiquitination of LRRK2 might target LRRK2 for degradation. To specifically measure changes in LRRK2 turnover *in vivo*, we created a doxycycline (“dox”)-inducible GFP-LRRK2 flow cytometric assay quantifying LRRK2 turnover: GFP-LRRK2 expression was induced to measurable but near physiologic levels (∼10 fold higher than endogenous LRRK2 expression), doxycycline removed, and GFP fluorescence measured by flow cytometry (schematic of experimental design is shown in Figure 3b). We first verified that normalized median GFP fluorescence intensity of GFP-LRRK2 was indeed proportional to LRRK2 levels on immunoblot (Figure S2c, S2d). We next tested the effect of TRIM1 expression on GFP-LRRK2 levels. As predicted, TRIM1 increased LRRK2 turnover—and thereby decreased LRRK2 levels (Figure 3c, quantified in 3d). LRRK2 levels did not change in the presence of TRIM1ΔRF, demonstrating TRIM1’s effect was E3-ubiquitin ligase-dependent (Figure 3d). We validated that the changes measured by GFP-fluorescence were reflected on immunoblot (Figure 3e, quantified in 3f). Consistent with our findings that TRIM1 C ubiquitinates LRRK2, LRRK2 levels were decreased by co-expression of TRIM1 C (Figure 3d). TRIM1 had no effect on turnover of dox-induced GFP alone (Figure S2e).

To define the pathway of TRIM1-catalyzed LRRK2 degradation, we measured TRIM1-mediated LRRK2 degradation in the presence of MG132 (proteasomal inhibitor) and chloroquine (autophagy inhibitor). LRRK2 degradation was inhibited by MG132, but not by chloroquine (Figure 3g), indicating that TRIM1-mediated degradation of LRRK2 occurs via the proteasome and not through autophagy. We validated by immunoblot that a second proteasomal inhibitor, bortezomib, restored TRIM1-mediated LRRK2 degradation (Figure 3h). Finally, we compared the effects of TRIM1 on LRRK2 levels to the effects of TRIM18 and CHIP in our dox-inducible GFP-LRRK2 line. Neither TRIM18 nor CHIP significantly decreased wild type GFP-LRRK2 steady-state levels in this assay (Figure S2f). Thus, TRIM1 is a microtubule-localized E3 ligase that ubiquitinates LRRK2, causing its degradation via the proteasome.

We performed ubiquitin-specific MS on immunoprecipitated LRRK2 to identify TRIM1-mediated polyubiquitin chain types and ubiquitination sites. In HEK-293T cells, FLAG-myc-LRRK2 and either GFP-TRIM1 WT, GFP-TRIM1 ΔRF, or GFP was expressed in the presence and absence of MG132. From each of these six conditions, LRRK2 was sequentially immunoprecipitated with anti-FLAG and anti-myc antibodies and then underwent ubiquitin-specific MS. K48 linkages (and no other ubiquitin linkage types) were identified in the LRRK2 sample containing WT GFP-TRIM1 and these K48 linkages were increased 3.5 fold in the presence of WT GFP-TRIM1 with MG132 (not shown). No polyubiquitin linkages were identified in samples containing GFP-TRIM1 ΔRF or GFP, with or without MG132. No ubiquitination sites were identified on LRRK2 in any of these six samples, including following enrichment for ubiquitinated peptides prior to MS. We therefore repeated the experiment in the presence of HA-ubiquitin and used sequential FLAG and HA immunoprecipitation to more specifically isolate ubiquitinated LRRK2. In this experiment, 92 of 176 lysine residues in LRRK2 were identified (60% sequence coverage of LRRK2). A single site of ubiquitination, LRRK2 K831 was identified and found to be dependent on TRIM1’s E3 ubiquitin ligase activity (Table S2, Figure S3a). However, LRRK2 K831R was still ubiquitinated by TRIM1 (Figure S3b), suggesting that TRIM1 ubiquitinates additional sites on LRRK2.

In this more sensitive experiment, we identified K48, K63, and K11 polyubiquitin linkages at ≥10-fold abundance in the presence of WT TRIM1 compared to the presence of TRIM1 ΔRF or control vector (Figure S3c). Using antibodies specific for K48 and K63 linkages, we identified TRIM1-mediated K48 but not K63 linkages on LRRK2 in the presence of HA-ubiquitin, which were also catalyzed by TRIM C but were not catalyzed TRIM1 ΔRF or TRIM18 (Figure S3d). To identify polyubiquitin chains directly conjugated to LRRK2 by TRIM1 in the absence of overexpressed ubiquitin, we used a pan-ubiquitin tandem ubiquitin binding entity (TUBE), which binds K6, K11, K48 and K63 polyubiquitin chains with nanomolar affinities. HEK-293T cells transfected with FLAG-LRRK2 and myc-TRIM1 or myc alone were treated with bortezomib and lysed in the presence of the deubiquitinase inhibitor PR-619. LRRK2 was immunoprecipitated with anti-FLAG antibodies, then eluted with FLAG peptide. From the LRRK2 elution, ubiquitinated LRRK2 was specifically isolated with a pan-selective TUBE (Figure S3e shows schematic of experiment). Lysates and eluates were immunoblotted with pan-ubiquitin, K48-specific, or K63-specific antibodies. TRIM1 increased the amount of total and K48-linked ubiquitin chains on LRRK2 but did not increase K63 linked chains (Figure S3f), demonstrating that TRIM1 mediates poly-K48 ubiquitination of LRRK2 to drive its proteasomal degradation. We were unable to identify a K11-chain specific antibody or TUBE and so have not completely ruled out that K11-linked polyubiquitin may also mediate LRRK2 proteasomal degradation by TRIM1.

### Knockdown of endogenous TRIM1 increases LRRK2 levels

To test the effect of endogenous TRIM1 on GFP-LRRK2 levels, we utilized a robust CRISPRi/dCas9 system^55^ to knockdown TRIM1 mRNA in conjunction with our flow cytometric GFP-LRRK2 assay. We generated dox-inducible GFP-LRRK2 cell lines stably expressing dCas9-BFP-KRAB. TRIM1 was knocked down using lentiviral transduction of sgRNA sequences targeted to the TRIM1 5’ UTR. TRIM1 sgRNA lowered TRIM1 mRNA levels to 20.0% +/- 4.4% compared to control sgRNA (Figure 4a) with a resulting increase in GFP-LRRK2 protein levels of 38.3% +/- 3.3% at 24 hours (Figure 4b). This increase in GFP-LRRK2 protein levels was significant throughout the length of the experiment (up to 44 hours post-dox withdrawal) (Figure S2g), indicating a persistent LRRK2 turnover deficit in these cells. Thus, knockdown of endogenous TRIM1 decreases turnover of GFP-LRRK2, consistent with an important role for TRIM1 in LRRK2 degradation.

**Figure 4.**
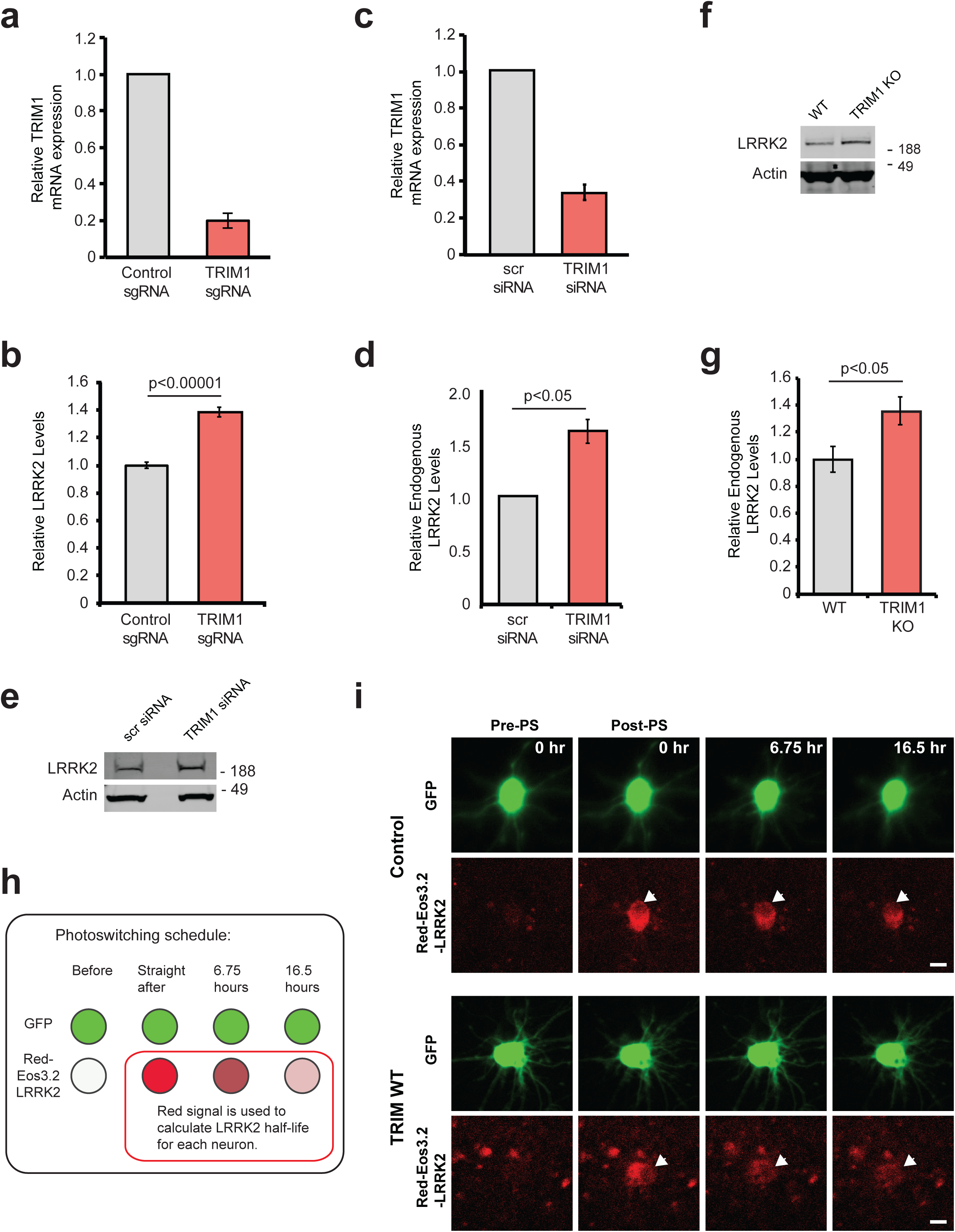
Reducing endogenous TRIM1 levels increases LRRK2 levels. (A) Relative TRIM1 mRNA expression in dox-inducible GFP-LRRK2 HEK-293T cells with dCas9 and either non-targeting sgRNA (grey bar) or four pooled TRIM1-targeting sgRNAs (red bar). (B) Quantification of GFP-LRRK2 fluorescence with TRIM1 knocked down (red bar) compared to cells with non-targeting sgRNA (grey bar) 24 hours after dox withdrawal. (C) Relative TRIM1 mRNA expression in Malme-3M cells with either scrambled siRNA (grey bar) or TRIM1-targeting siRNA (red bar). (D) Quantification of endogenous LRRK2 levels measured by immunoblots following TRIM1 siRNA knock down showing mean value from six independent experiments with error bars showing standard error of the mean. (E) Representative immunoblot of endogenous LRRK2 in lysate of Malme-3M cells with scrambled siRNA (left lane) or endogenous TRIM1 knocked down by targeted siRNA (right lane). (F) Immunoblot of endogenous LRRK2 in WT and TRIM1 KO HEK-293T cells. (G) Quantification of (F) showing mean value from four independent experiments with error bars showing standard error of the mean. (H) Schematic of optical pulse labeling experiment in which primary cortical neurons were co-transfected with mEos3.2-LRRK2 and GFP, and either TRIM1 or control plasmid. Cells were pulsed for 5-8 s with 405-nm light, causing a portion of mEos3.2-LRRK2 to fluoresce red, and cells were imaged over the indicated time period. (I) Representative primary cortical neurons from photoswitching experiment. Prior to photoswitching (Pre-PS), mEos3.2-Red-LRRK2 is not detected. After photoswitching (Post-PS) mEos3.2-Red-LRRK2 is detected, nuclear excluded (arrow), and decays with time. LRRK2 decays faster in neurons transfected with TRIM1. Scale bar is 10 μM. P-value for Panel B calculated using a t-test. Significance testing for panels D and G calculated was performed using Mann-Whitney U test.

To examine the effects of TRIM1 on endogenous LRRK2 levels, we utilized human melanoma Malme-3M cells, which express relatively high levels of both LRRK2 and TRIM1 mRNA (NCBI Geoprofiles ID#86805339 and #86784306). SiRNA against TRIM1 was used to knockdown TRIM1 mRNA levels to 33% +/- 6% relative to siRNA scrambled control (Figure 4c). TRIM1 knockdown resulted in an almost 2-fold increase (162% +/- 13%) in endogenous LRRK2 levels at 48 hours (Figures 4d and 4e). We also measured LRRK2 levels in TRIM1 CRISPR knockout versus WT HEK-293T cells. In HEK-293T cells, the TRIM1 KO showed increased LRRK2 levels at steady state (136% +/- 8% in TRIM1 KO vs 100% +/- 7% in WT, Figures 4f and 4g). Thus, TRIM1 is a key regulator of endogenous LRRK2 turnover.

### TRIM1 mediates LRRK2 turnover in neurons

We next tested TRIM1’s ability to drive LRRK2 turnover in primary cortical neurons using optical pulse labeling, a method that has been used to monitor turnover of several neurodegenerative proteins, including huntingtin^56^, α-synuclein^57^ and TDP-43.^58^ In this assay, we quantified LRRK2 protein level within individual neurons over multiple time points using the photoswitchable protein mEos3.2 fused to LRRK2. Cells expressing mEos3.2-LRRK2 initially fluoresce green; however, upon illumination with a 405 nm wavelength light, a population of green-mEos3.2-LRRK2 is irreversibly switched to red, creating a distinct pool of red-mEos3.2-LRRK2 in each neuron. Individual neurons were followed over time, and red-mEos3.2-LRRK2 signal was quantified using automated longitudinal imaging, allowing us to derive individual LRRK2 half-life measurements for each neuron (see schematic in Figure 4h). Embryonic day 20-21 rat primary cortical neurons were co-transfected with pGW1-GFP as a morphology marker, pGW1-mEos3.2- LRRK2 and either TRIM1 or a control plasmid, and photoswitched with a 5-8-second pulse of light at 405-nm wavelength. Neurons were imaged every 4-10 hours for red and green fluorescence using custom-based automated algorithms to capture images in an unbiased and high-throughput manner. Representative neurons in the presence and absence of TRIM1 are shown in Figure 4i. In this neuronal system, as in the HEK-293T cell system, LRRK2 decay was significantly accelerated by almost two-fold in the presence of TRIM1 (t_1/2_ LRRK2 = 24.9 h in the absence of exogenous TRIM1, t_1/2_ LRRK2 = 15.9 h in the presence of exogenous TRIM1, p =0.025, 113 and 87 neurons per group respectively, 3 independent experiments).

### The interdomain region between LRRK’s ankyrin and LRR domains binds TRIM1

In order to better define the TRIM1/LRRK2 interaction, we performed a series of co-immunoprecipitation experiments using truncation mutants of both proteins (domain structure of LRRK2 truncation mutants illustrated in Figure S4a). We found that LRRK2 interacts with the tandem B-box domain of TRIM1 (Figure 5a) with binding most dependent on TRIM1’s linker and B-box1 domain (Figure 5b). Notably, with the exception of the extreme C-terminus, this domain includes the portion of least homology to TRIM18 (Figure S1d, double red line), suggesting that variations in this region may account for the differential ability of these highly homologous TRIM family members to bind LRRK2.

**Figure 5.**
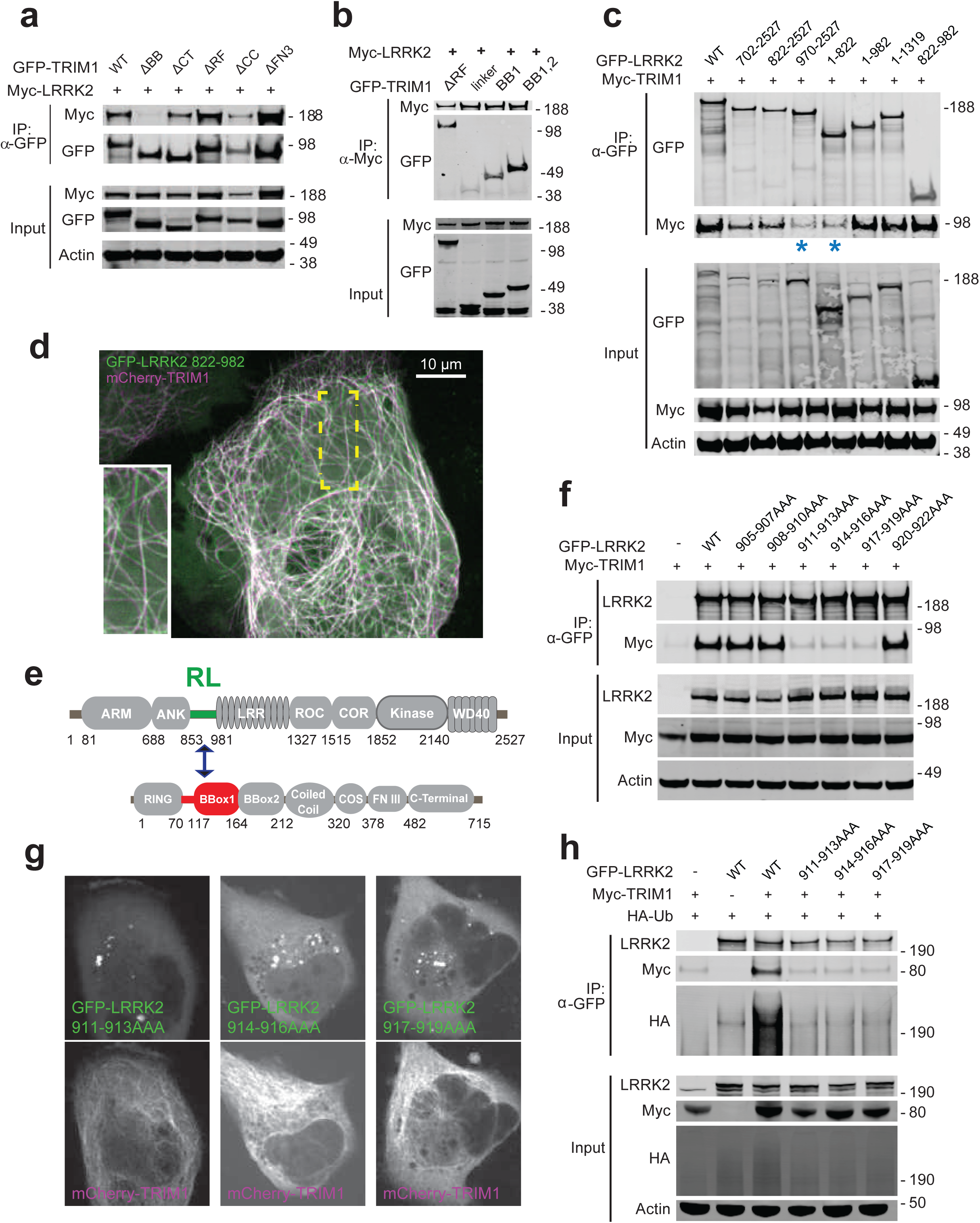
TRIM1 binds an N-terminal LRRK2 regulatory loop region via its B-box domain. (A) Co-immunoprecipitation of full-length myc-LRRK2 with GFP-TRIM1 domain constructs in HEK-293T cells (ΔBB: TRIM1 construct lacking both B-box domains; ΔCT: TRIM1 lacking C-terminal domain, ΔRF: TRIM1 lacking ring-finger domain; ΔCC: TRIM1 lacking coiled coil domain; ΔFN3: TRIM1 lacking fibronectin III domain; details of constructs in ^74^. (B) Co-immunoprecipitation of full-length myc-LRRK2 with GFP-TRIM1 B-box domain constructs in HEK-293T cells (ΔRF denotes TRIM1_70-715_; linker denotes TRIM1_70-117_; BB1 denotes TRIM1_70-164_; BB1,2 denotes TRIM1_70-212_). (C) Co-immunoprecipitation of full-length myc-TRIM1 with GFP-LRRK2 domain constructs in HEK-293T cells. LRRK2 constructs indicate included amino acids from full length LRRK2 sequence and are illustrated in Figure S4. LRRK2_822-892_ is sufficient for interaction with TRIM1. (D) Live-cell confocal microscopy of GFP-LRRK2_822-982_ and mCherry-TRIM1 transiently transfected into H1299 cells. Inset shows higher magnification of region identified by the yellow box. (E) Schematic of LRRK2-TRIM1 domain interaction mediated by the LRRK2 Regulatory Loop (“RL,” green) and TRIM1_BBox1_ (red). (F) Co-immunoprecipitation of full-length myc-TRIM1 with GFP-LRRK2 WT and RL alanine scanning mutants. Mutants are full length LRRK2 constructs with the three amino acids residues indicated mutated to three alanines. (G) Live-cell confocal microscopy of GFP-LRRK2 RL alanine scanning mutants and mCherry-TRIM1 transiently transfected into H1299 cells. (H) Ubiquitination of immunoprecipitated GFP-LRRK2 WT versus RL alanine scanning mutants. All co-immunoprecipitation and microscopy experiments are a representative image of at least three independent experiments.

LRRK2 constructs lacking the interdomain region between the ankyrin and LRR domains (amino acids 822-982) were markedly reduced in their ability to bind full-length TRIM1 (Figure 5c, asterisks denote constructs with strongly decreased binding, and S4a). A truncated LRRK2_822-982_ mutant was sufficient for binding to TRIM1 (Figure 5c, far right lane). LRRK2_822-982_ was also sufficient for TRIM1-mediated LRRK2 localization to microtubules (Figures 5d and S4a). Interestingly, the interdomain region that binds TRIM1 is absent from LRRK2’s closest homologue, LRRK1.^59^ This region is already known to be critical in mediating binding of 14-3-3 proteins, LRRK2’s best understood interactors.^13^ It has also been shown to undergo significant phosphorylation in response to upstream kinases, suggesting it is a key LRRK2 regulatory region.^33^ *In silico* modelling of the secondary structure of LRRK2_822-982_ predicts it to be >75% unstructured and >75% solvent exposed (www.predictprotein.org), which is consistent with very recent cryo-EM structure of full-length LRRK2, which demonstrated a hinge helix (amino acids 834-852) followed by an unstructured region not amenable to cryoEM (amino acids 853-981).^31^ From hereon, we designate LRRK2_853-981_ the LRRK2 Regulatory Loop (LRRK2 RL) region (Figure 5e). Within LRRK2 RL, we performed alanine scanning to pinpoint the precise amino acids required for LRRK2’s interaction with TRIM1. We identified a 9 amino acid region (amino acids 911-919) required for LRRK2 binding as measured by co- immunoprecipitation (Figure 5f). Consistently, these 9 amino acids were also required for TRIM1 to cause LRRK2 microtubule localization (Figure 5g) and for TRIM1 to ubiquitinate LRRK2 (Figure 5h).

### LRRK2 RL phosphorylation influences TRIM1 versus 14-3-3 binding

Interaction of 14-3-3 with LRRK2 has been studied in detail.^32^ This interaction depends on the phosphorylation state of multiple LRRK2 serine residues, with Ser910 and Ser935 phosphorylation absolutely required for the LRRK2-14-3-3 interaction.^60^ Since Ser910 and Ser935 are located within LRRK2 RL, directly adjacent to the 9 amino acids required for TRIM1 binding, we postulated that TRIM1 and 14-3-3 might compete for LRRK2 binding. Specifically, we hypothesized that Ser910 and Ser935 phosphorylation might serve as a molecular switch controlling LRRK2 RL’s predilection for binding partners. To visualize the subcellular localization of LRRK2 in the presence of both 14-3-3 and TRIM1, we transfected H1299 cells with mCherry-LRRK2, EBFP2-14-3-3 theta, and GFP-TRIM1. In the absence of GFP-TRIM1, both mCherry-LRRK2 and EBFP2-14-3-3 showed a diffusely cytoplasmic localization (Figure S4b). In the presence of GFP-TRIM1, mCherry-LRRK2 associated with microtubules, while EBFP2-14-3-3 remained diffusely cytoplasmic (Figure 6a and S4c). We quantified the proportion of cells with microtubule-associated mCherry-LRRK2 in the presence of EBFP2-14-3-3 and/or GFP-TRIM1 and found that TRIM1 caused LRRK2 localization to microtubules in essentially all cells in which both proteins were expressed, regardless of the presence of overexpressed 14-3-3 (94.2% +/- 1.7% without 14-3-3; 92.0% +/- 2.6% with 14-3-3, mean +/- S.D.; Figure 6b). No cells showed mCherry-LRRK2 at microtubules in the presence of EBFP2-14-3-3 alone (0% +/- 0%, mean +/- S.D.). Thus, GFP-TRIM1 causes mCherry-LRRK2 microtubule association in both the absence and presence of overexpressed EBFP2-14-3-3.

**Figure 6.**
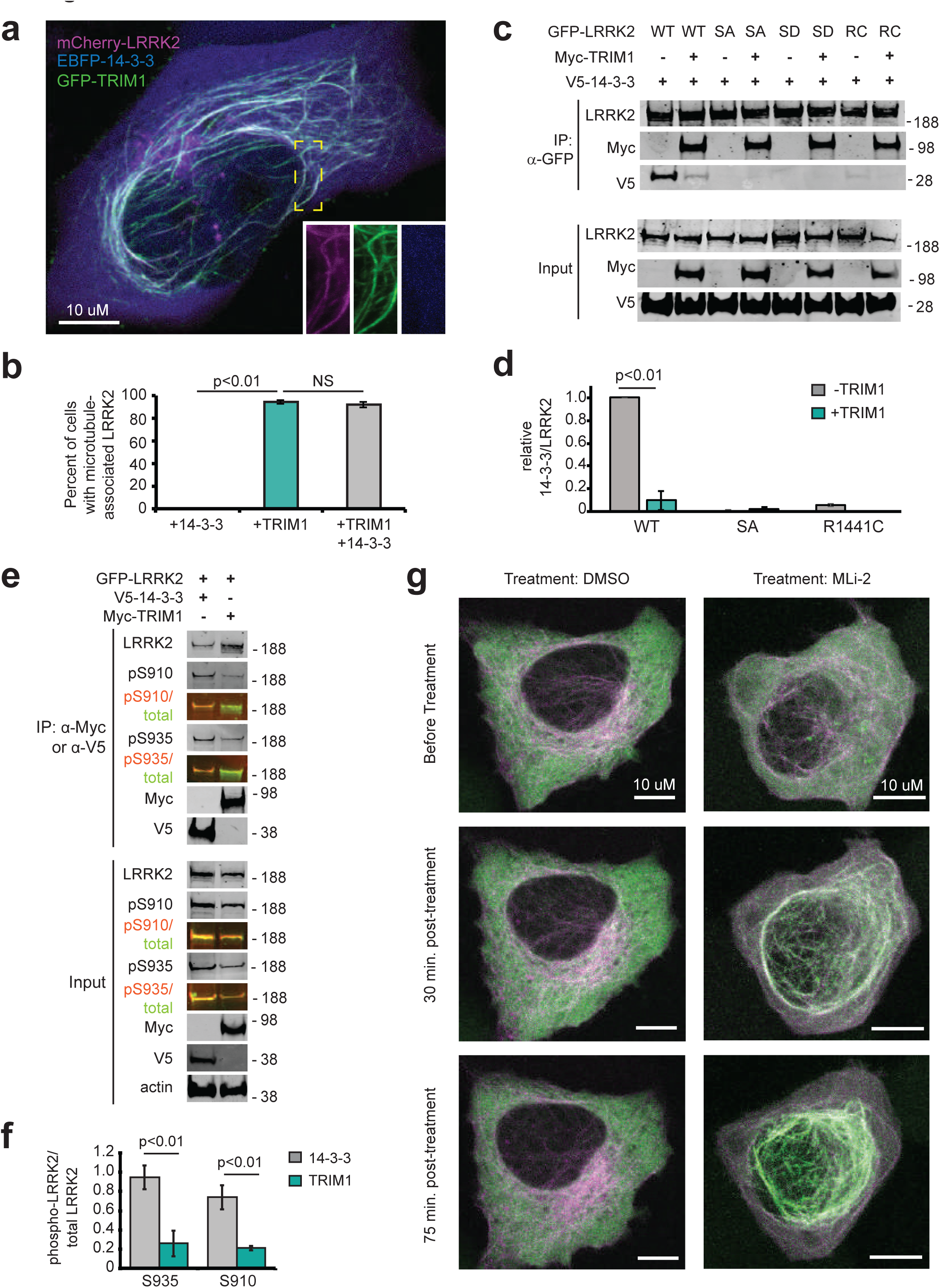
TRIM1 competes with 14-3-3 to bind LRRK2’s regulatory loop and recruit LRRK2 to microtubules. (A) Live-cell confocal microscopy of mCherry-LRRK2 in the presence of EBFP2-14-3-3 and GFP-TRIM1. (B) Quantification of H1299 cells with microtubule-associated LRRK2 when co-expressed with indicated proteins. 100 cells were evaluated in each experiment, bars indicate mean +/- standard deviation (two independent experiments). (C) Co-immunoprecipitation of GFP-LRRK2 wild type (WT), Ser910Ala Ser935Ala (SA), Ser910Asp Ser935Asp (SD), or R1441C (RC) with V5-14-3-3 theta in the presence and absence of myc-TRIM1 in HEK-293T cells. (D) Quantification of (C) showing mean value from three independent experiments with error bars showing standard deviation. Significance determined by Mann-Whitney U test (E) Co-immunoprecipitation of GFP-LRRK2 with either V5-14-3-3 theta or myc-TRIM1 in HEK-293T cells. Overlaid immunoblots in color show relative ratio of total to phospho-LRRK2 (total LRRK2 in green, antibody is NeuroMab clone N241A/34; phospho-LRRK2 is in red, antibodies are phospho-Ser910 (Abcam, UDD 1 15(3)) and phospho-Ser935 (Abcam, UDD 2 10(12)). (F) Quantification of (E) with mean +/- standard error of the mean from three independent experiments. (G) Live-cell confocal microscopy of GFP-LRRK2 in the presence of mCherry-TRIM1 following treatment with LRRK2 kinase inhibitor MLi-2 (200 nM) or vehicle. Rare cells with low levels of co-localization prior to treatment were followed. LRRK2 is shown in green and TRIM1 in purple. Images from isolated channels are shown in Figure S5C. All live cell images and co- immunoprecipitation experiments are a representative image of at least three independent experiments. Significance testing for panel B was performed using Kruskal-Wallis with post-hoc Dunn test and Bonferroni correction and Mann-Whitney U test for panels D and F.

To characterize residues in LRRK2 that influence its binding to TRIM1 vs. 14-3-3, we quantified co-immunoprecipitation of 14-3-3 with various LRRK2 point mutants in the presence and absence of TRIM1. GFP-LRRK2-WT and V5-14-3-3 theta co- immunoprecipitated in the absence of myc-TRIM1, as reported in the literature (Figure 6c).^13, 61^ In the presence of both V5-14-3-3 and myc-TRIM1, GFP-LRRK2-WT robustly co-immunoprecipitated myc-TRIM1 but only bound 19% as much V5-14-3-3 as it did in the absence of myc-TRIM1 (Figure 6d), demonstrating that TRIM1 can disrupt LRRK2’s binding to 14-3-3. To test whether Ser910 and Ser935 phosphorylation is required for LRRK2 binding to TRIM1, we mutated GFP-LRRK2 Ser910 and Ser935 to non-phosphorylatable alanines (“GFP-LRRK2 SA”). As demonstrated by others, GFP-LRRK2-SA did not bind V5-14-3-3.^13^ In contrast, myc-TRIM1 co-immunoprecipitated GFP-LRRK2-SA to the same extent as GFP-LRRK2-WT (Figure 6c). Lack of binding of GFP-LRRK2-SA to 14-3-3 did not change in the presence of myc-TRIM1 (Figure 6d). Thus, TRIM1 strongly binds both LRRK2 WT and non-phosphorylatable LRRK2-SA, while 14- 3-3 selectively binds LRRK2 WT, in which S910 and S935 can be phosphorylated. Similarly, the LRRK2 PD mutant LRRK2 R1441C, which lacks S910 and S935 phosphorylation and does not bind 14-3-3, still strongly bound myc-TRIM1 but did not co-immunoprecipitate with V5-14-3-3 (Figures 6c and d). Finally, we attempted to construct a phosphomimetic version of GFP-LRRK2 by mutating Ser910 and Ser935 to aspartic acid (GFP-LRRK2 SD). TRIM1 bound GFP-LRRK2-SD to a similar extent to GFP-LRRK2-WT. However, GFP-LRRK2-SD did not bind 14-3-3 in the presence or absence of TRIM1, indicating that GFP-LRRK2-SD does not adequately mimic phosphorylated Ser910 and Ser935 (Figure 6c). Thus this construct does not provide information regarding the extent to which phosphorylated LRRK2 binds TRIM1.

We subsequently measured the phosphorylation state of LRRK2 bound to either TRIM1 or 14-3-3 using co-immunoprecipitation followed by quantitative immunoblot with phospho-specific antibodies against either phospho-Ser910 or phospho-Ser935 LRRK2. LRRK2 bound to 14-3-3 showed markedly increased phosphorylation of both Ser910 and Ser935 compared to LRRK2 bound to TRIM1 (Figure 6e, compare red (phospho) to green (total) signal of immunoprecipitated LRRK2). We quantified the ratio of phospho-LRRK2 to total LRRK2 signal for 14-3-3-bound LRRK2 and TRIM1-bound LRRK2 (normalized to phospho-LRRK2:total-LRRK2 in the input lysate) (Figure 6f). Ser935 phosphorylation of TRIM1-bound LRRK2 was 27% of 14-3-3-bound LRRK2 (ratio of phospho-S935 LRRK2:total LRRK2 signal was 0.26 +/- 0.13 for TRIM1-bound LRRK2 and 0.94 +/- 0.13 for 14-3-3-bound LRRK2, mean +/- S.D.). Similarly, LRRK2 Ser910 phosphorylation of TRIM1-bound LRRK2 was 28% of 14-3-3-bound LRRK2 (ratio of phospho-S910 LRRK2: total LRRK2 was 0.21 +/- 0.02 for TRIM1-bound LRRK2 and 0.74 +/- 0.12 for 14-3-3- bound LRRK2, mean +/- S.D.). Thus, phosphorylation of LRRK2’s RL region influences LRRK2’s affinity for partner proteins, with a larger proportion of unphosphorylated LRRK2 bound to TRIM1 than to 14-3-3. Together, these data suggest that TRIM1 preferentially binds non-phosphorylated LRRK2 and localizes it to microtubules while 14-3-3 preferentially binds phosphorylated LRRK2 in the cytosol.

### Type 1 LRRK2 kinase inhibitors increase TRIM1-LRRK2 association

Type 1 LRRK2 kinase inhibitors such as MLi-2 increase LRRK2’s microtubule localization and cause LRRK2 ubiquitination and proteasomal degradation through unknown mechanisms.^41, 62^ Because TRIM1 expression phenocopies the effects of MLi-2 treatment (i.e., causes LRRK2 localization to microtubules and leads to LRRK2 ubiquitination and degradation), we hypothesized that TRIM1 may be required for LRRK2 degradation following Type 1 inhibitor treatment. We first tested whether MLi-2 treatment increases colocalization of LRRK2 and TRIM1 at microtubules. H1299 cells transfected with GFP-LRRK2 and mCherry-TRIM1 were treated with MLi-2 and time-lapse live cell microscopy was performed. We focused on the rare subset of cells with predominantly cytoplasmic GFP-LRRK2 in the presence of mCherry-TRIM1 prior to treatment with MLi-2. Thirty minutes post-treatment with 200 nM MLi-2, GFP-LRRK2 and mCherry-TRIM1 were strongly associated at microtubules (Figure 6g, S5a), which continued for the duration of the experiment (120 minutes). GFP-LRRK2 association with TRIM1 and with the microtubule network did not increase after vehicle-alone treatment. To test the effect of MLi-2 on association of endogenous LRRK2 and TRIM1, we co-immunoprecipitated LRRK2 with TRIM1 in HEK-293T cells treated with MLi-2 or vehicle control. Under endogenous conditions, we observed increased association of LRRK2 and TRIM1 in the presence of MLi-2 compared to the absence of MLi-2 (Figure 1e, lanes 2 and 4).

To test whether TRIM1 is required for LRRK2 degradation following Type 1 kinase inhibitor treatment, we measured GFP-LRRK2 levels after MLi-2 or vehicle treatment using the dCas9/dox-GFP-LRRK2 system. MLi-2 treatment at 100 nM for 24 hours decreased GFP-LRRK2 levels by about 50%, consistent with what others have observed (Figure S5b, compare vehicle to MLi-2 treatment in the presence of control sgRNA).^18, 62^ If TRIM1 is required for LRRK2 degradation following MLi-2 treatment, TRIM1 knockdown should rescue LRRK2 levels in the presence of MLi-2. We therefore measured the effect of TRIM1 knockdown on LRRK2 levels following MLi-2 treatment. As we had previously shown (see Figures 4b, S2g), in the absence of MLi-2, TRIM1 knockdown increased LRRK2 levels compared to control sgRNA. In the presence of MLi-2, TRIM1 knockdown caused no additional rescue of LRRK2 levels compared to control sgRNA (Figure S5b; vehicle-treated LRRK2 levels are normalized to one). In control cells, MLi-2 treatment led to a 52.2% +/- 3% decrease in total LRRK2 levels compared to vehicle, and in cells with TRIM1 knocked down, MLi-2 caused a 54.8% +/- 4.4% decrease in LRRK2 compared to vehicle. We thus conclude that while TRIM1 mediates basal LRRK2 turnover, TRIM1 is not required for LRRK2 degradation following MLi-2 treatment. Hence, the mechanisms responsible for kinase inhibitor-mediated LRRK2 turnover remain to be discovered.

### TRIM1 inhibits LRRK2 kinase activation by Rab29

Rab29, which is found at Golgi network membranes, was recently identified as a strong activator of LRRK2 kinase function in cell-based overexpression systems.^24, 29^ Rab29 overexpression increases LRRK2 autophosphorylation at Ser1292 and LRRK2 phosphorylation of substrate Rab proteins (Rab10 at Thr73 and Rab29 at Thr71; these phosphorylation sites are LRRK2-specific).^24^ The N-terminal portion of LRRK2 interacts with Rab29. In particular, the C-terminal half of LRRK2’s armadillo domain is critical for LRRK2-Rab29 interaction.^30^ Conserved Leu-rich motifs in LRRK2’s ankyrin domain— which is near LRRK2 RL—also appear essential for LRRK2 activation by Rab29.^24^ We hypothesized that TRIM1 might inhibit the ability of Rab29 to activate LRRK2’s kinase activity. To measure TRIM1’s effect on Rab29-mediated LRRK2 activation, we dox-induced GFP-LRRK2 WT or R1441G expression in HEK-293T cells and co-expressed myc-TRIM1 and/or HA-Rab29 via transient transfection. In these experiments, we provided continuous high-level (1 µg/ml) dox-induction until the time of harvest to maintain equivalent LRRK2 levels in the presence and absence of TRIM1 and used quantitative immunoblot to validate that total LRRK2 levels were equivalent in all conditions (not shown).

Consistent with others’ work, overexpressed Rab29 increased LRRK2 WT and LRRK2 R1441G kinase activity ∼2-4 fold, as measured by autophosphorylation of LRRK2 Ser1292 (Figure 7a, quantified in 7b).^24^ Coexpression of TRIM1 in the setting of Rab29 overexpression decreased phosphorylation of LRRK2 Ser1292 to baseline levels (i.e., levels observed without Rab29 overexpression) for both LRRK2 WT and LRRK2 R1441G. TRIM1 coexpression had no effect on Ser1292 phosphorylation in the absence of Rab29 (Figures 7a and b). We quantified phosphorylation of Rab29 at Thr71 and Rab10 at Thr73 as additional measures of LRRK2 kinase activity. Coexpression of TRIM1 decreased phosphorylation of Rab29 Thr71 by about half for LRRK2 WT (44% +/- 5%, mean +/- S.D.) and LRRK2 R1441G (58% +/- 7%, mean +/- S.D.) (Figure 7a, quantified in 7c). Co-expression of TRIM1 modestly decreased phosphorylation of endogenous Rab10 Thr73 by LRRK2 WT (70% +/- 9%, mean +/- S.D.) and LRRK2 R1441G (71% +/- 8%, mean +/- S.D.), but did not fully restore to baseline levels without Rab29 (Figure 7a, quantified in 7d). Importantly, co-expression of only TRIM1 with LRRK2 had no effect on Thr73 Rab10 phosphorylation, similar to our findings for LRRK2 Ser1292. Together, these data show that TRIM1 inhibits Rab29-mediated LRRK2 kinase activation but does not appear to have an effect on basal LRRK2 kinase activity.

**Figure 7.**
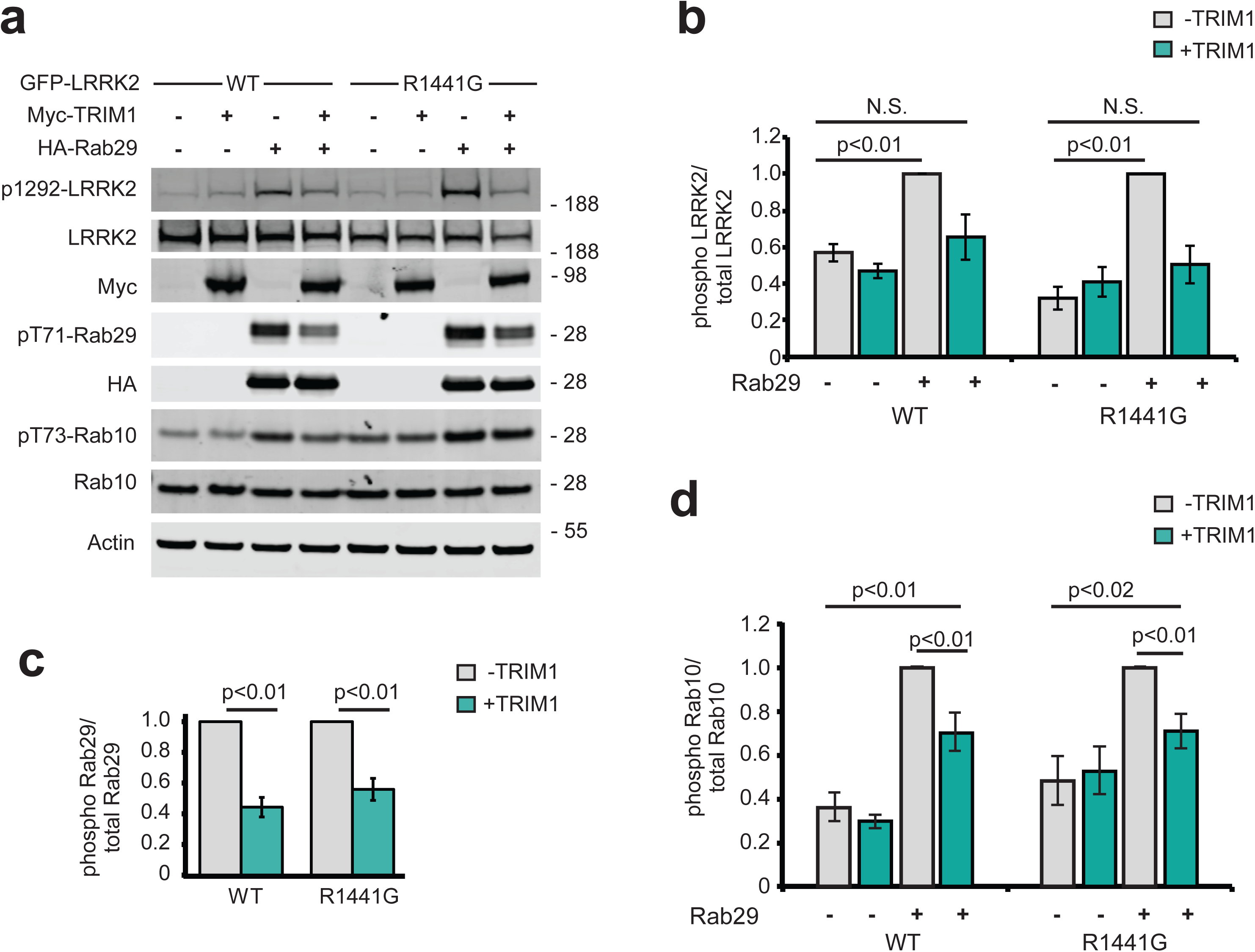
TRIM1 decreases LRRK2’s activation by Rab29. (A) Immunoblot of LRRK2 phosphorylation at Ser1292, Rab29 phosphorylation at Thr71, and Rab10 phosphorylation at Thr73 in the presence and absence of TRIM1 for wild type LRRK2 and LRRK2-PD mutant R1441G. (B) Quantification of LRRK2 autophosphorylation in (A). (C) Quantification of Rab29 Thr71 phosphorylation in (A). (D) Quantification of Rab10 Thr73 phosphorylation in (A). Quantifications show the mean value from 3-4 independent replicates with error bars showing standard error of the mean. Significance testing for panels B-D was performed using Kruskal-Wallis with post-hoc Dunn and test Bonferroni correction.

To begin to dissect the mechanism by which TRIM1 inhibits Rab29-mediated LRRK2 kinase activation, we measured phosphorylation of Rab29 Thr71 in the presence of WT TRIM1, TRIM1 C (intact E3 ligase activity and cytoplasmic), TRIM1 ΔRF (no E3 ligase enzymatic activity but still binds microtubules), or control vector. We utilized phosphorylation of Rab29 Thr71 because it is our most robust readout of LRRK2 kinase activation: on quantitative immunoblot, phospho-Thr71 Rab29 signal is 20-100x higher than phospho-Ser1292 LRRK2 or phospho-Thr73 Rab10 (see Figure 7a). Identical to WT TRIM1, TRIM1 C inhibited Rab29-mediated LRRK2 activation for both LRRK2 WT and R1441G (Figure 8a, quantified in Figure 8c top panel). TRIM1 ΔRF did not inhibit Rab29-mediated LRRK2 activation, suggesting that TRIM1’s E3 ubiquitin ligase domain is required for inhibition. Mutation of Lys831, the ubiquitinated LRRK2 residue identified by MS, to Arg (LRRK2 K831R) did not rescue TRIM1’s inhibition of Rab29-mediated LRRK2 activation (Figure S5c, quantified in S5d).

**Figure 8.**
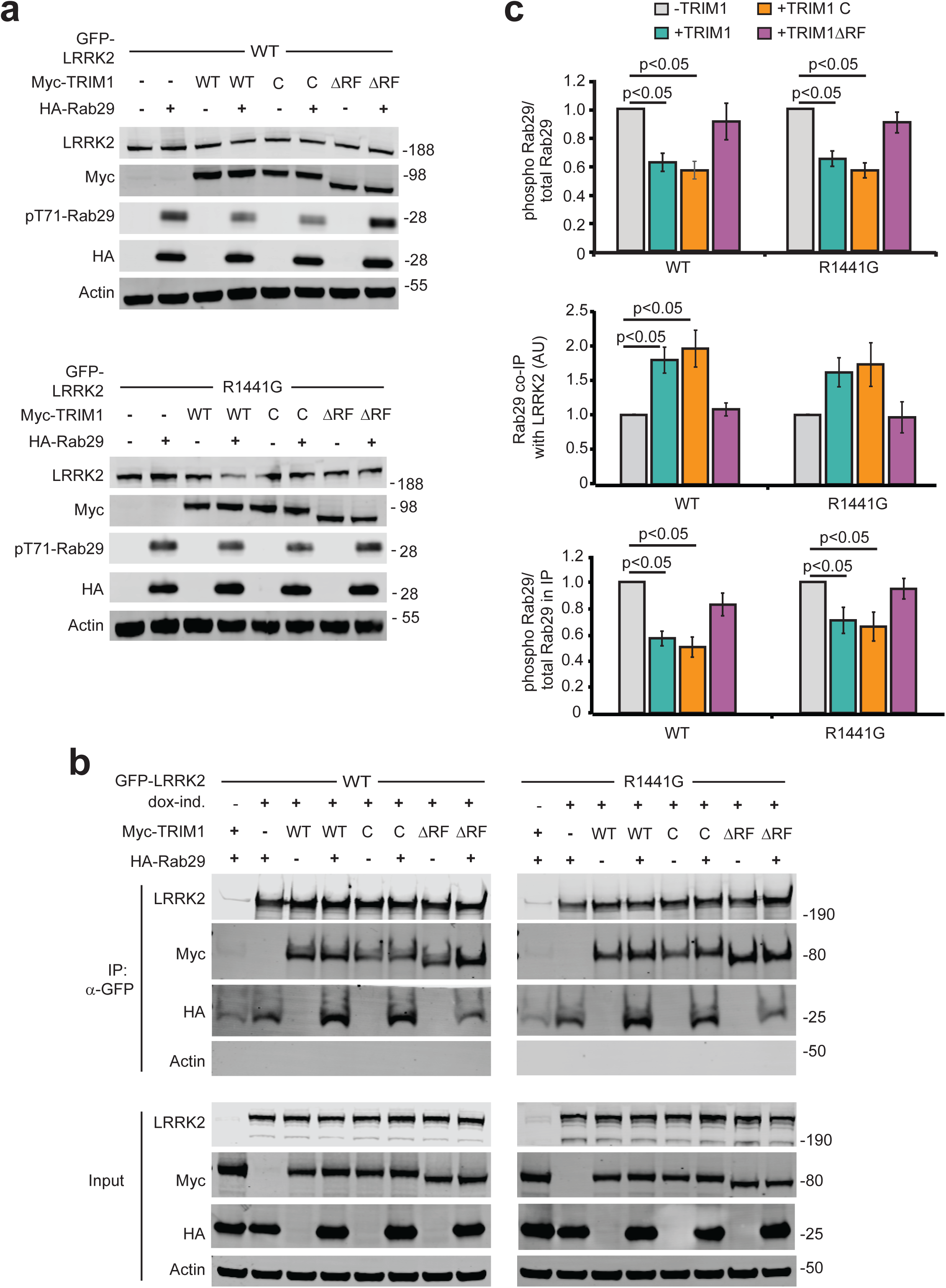
The LRRK2-Rab29 interaction is modulated by TRIM1’s E3 ligase activity. (A) Immunoblot of Rab29 phosphorylation with wild type LRRK2 (top) or LRRK2-PD mutant R1441G (bottom) in the absence of overexpressed TRIM1 or with overexpression of myc-tagged wild type TRIM1 (WT), microtubule-nonbinding TRIM1 (TRIM1 C), or TRIM1 lacking E3-ligase function (TRIM1ΔRF). (B) Co-immunoprecipitation of Rab29 and TRIM1 variants with LRRK2 WT or R1441G. (C) Quantification of: Rab29 phosphorylation relative to total Rab29 in lysate from part A (top panel), Rab29 co-IPed with LRRK2 from part B (middle panel) and Rab29 phosphorylation relative to total Rab29 in co-IP with LRRK2 (bottom panel). All immunoblots are representative images and quantification shows the mean value from at least three independent experiments with error bars showing the standard error. Significance testing for panel C was performed using Kruskal-Wallis with post-hoc Dunn test and Bonferroni correction.

We further investigated the mechanism of TRIM1’s inhibition of Rab29-mediated LRRK2 kinase activation using co-immunoprecipitation and quantitative immunoblotting to measure the interaction of LRRK2 and Rab29 in the presence of TRIM1 constructs (WT, C, ΔRF) or control vector. Intriguingly, TRIM1 constructs with retained E3 ligase function (WT, C) increased co-immunoprecipitation of Rab29 with WT LRRK2 relative to control vector while TRIM1 ΔRF did not (Figure 8b, quantified in 8c middle panel). LRRK2 R1441G showed the same trend, though the findings did not reach significance using non-parametric multiple comparison testing. Interestingly, Rab29 co-immunoprecipitated with LRRK2 showed the same pattern of Thr71 phosphorylation as total (lysate) Rab29, with the fraction of phosphorylated Rab29 decreased in the presence of TRIM1 WT and TRIM1 C but not in the presence of TRIM1 RF (Figure 8c bottom panel). While further work is required to delineate the precise mechanisms by which TRIM1 can inhibit Rab29-mediated LRRK2 activation, these findings suggest that E3-ligase active TRIM1 can modulate the interaction of Rab29 and LRRK2. One possible mechanism is that ubiquitination changes the binding properties of LRRK2 to Rab29, increasing association of these proteins.

### TRIM1 rescues the neurite outgrowth defect caused by LRRK2 G2019S

The most common LRRK2 PD-driving point mutation is LRRK2 G2019S. LRRK2 G2019S expression is known to cause decreased neurite outgrowth, a microtubule-driven process that reflects neuronal health.^63, 64^ Similar to wild type LRRK2, LRRK2 G2019S was drawn to microtubules in H1299 cells (Figure 9a) and ubiquitinated (Figure 9b) and degraded via the proteasome (Figure 9c) in HEK-293T cells in a TRIM1-dependent manner. To test if TRIM1 rescues LRRK2 G2019S-driven neurite outgrowth deficits, a classic cellular model of LRRK2-PD toxicity, we used a previously published rat PC12 pheochromocytoma cell line harboring dox-inducible LRRK2 G2019S (“PC12 dox-LRRK2 G2019S”).^65^ We validated that type 1 LRRK2 kinase inhibitors rescue the neurite outgrowth deficiency caused by induction of LRRK2 G2019S in this cell line, demonstrating the phenotype is LRRK2-dependent (not shown) and that TRIM1 caused microtubule localization of GFP-LRRK2 G2019S (Figure S5e) and LRRK2 ubiquitination in PC12 cells (Figure S5f).

**Figure 9.**
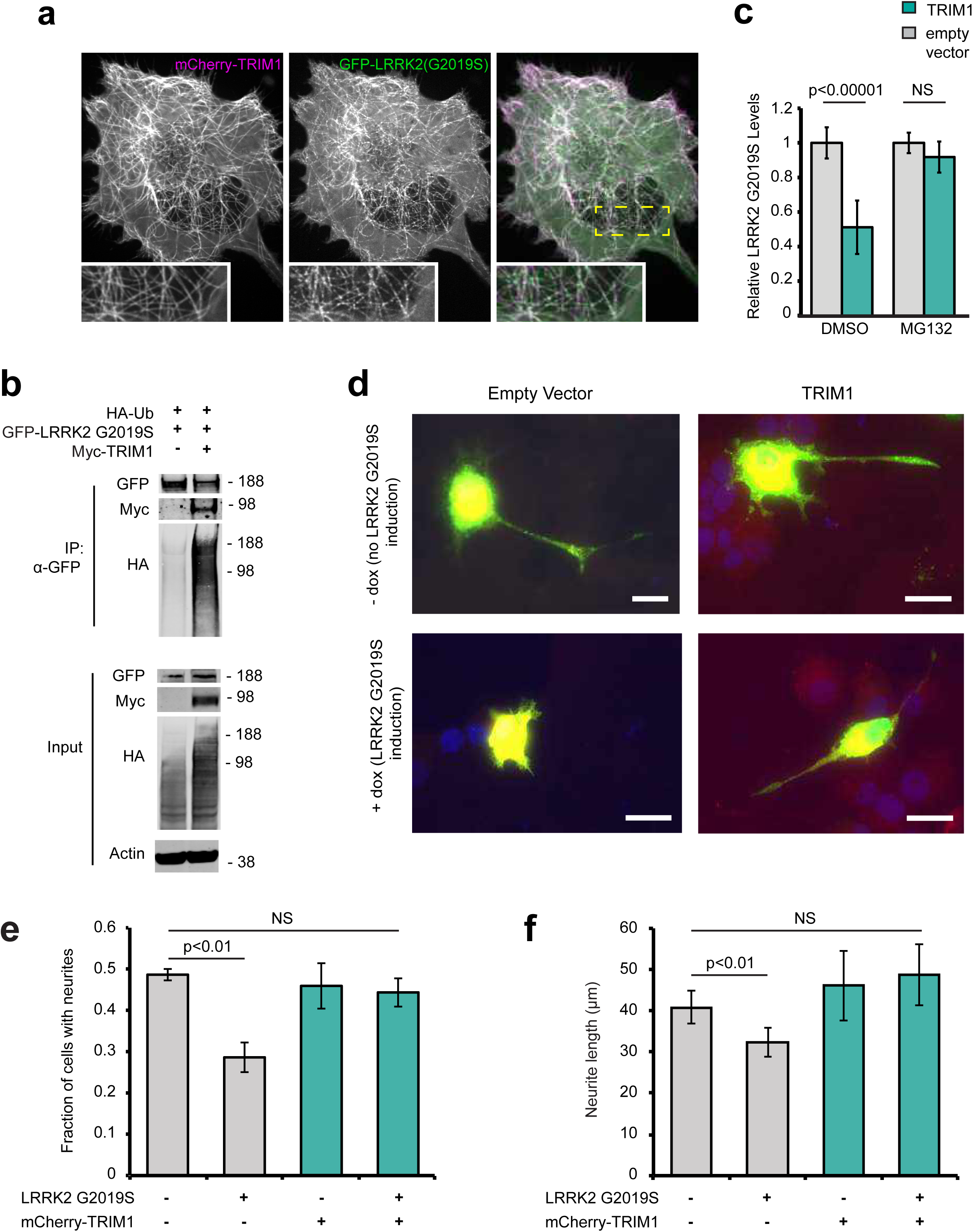
TRIM1 mediates proteasomal degradation of PD-mutant LRRK2 G2019S to rescue its toxicity. (A) Live-cell confocal microscopy of GFP-LRRK2 G2019S and mCherry-TRIM1 transiently transfected into H1299 cells. (B) Co-immunoprecipitation and ubiquitination of GFP-LRRK2 G2019S with myc-TRIM1 in the presence of HA-ubiquitin in HEK-293T cells. (C) Flow cytometric assay on dox-inducible GFP-LRRK2 G2019S HEK-293T cells in the presence and absence of TRIM1 and the proteasome inhibitor MG132; bars show median green fluorescence intensity with error bars showing twice the standard error of the mean. (D) Representative dox-inducible LRRK2 G2019S PC-12 cells transfected with mCherry-TRIM1 or mCherry alone vector and GFP and differentiated with NGF for 5 days in the presence and absence of 1 µg/ml doxycycline. (E) Quantification of the fraction of neurite-bearing PC-12 cells in the presence and absence of LRRK2 G2019S and the presence and absence of TRIM1; bars show average of three independent experiments of 150-250 cells each, error bars show standard error of the mean. (F) Quantification of average neurite length on PC-12 cells with neurites in the presence and absence of LRRK2 G2019S and the presence and absence of TRIM1; bars show average of three independent experiments, error bars show standard error of the mean. Significance testing for panel C was performed using ANOVA with post-hoc t-test with Bonferroni correction. Significance testing for panel E was performed using a test for equality of binomial parameters and F was performed using Kruskal-Wallis with post-hoc Dunn test and Bonferroni correction.

PC12 dox-LRRK2 G2019S cells were transfected with mCherry-TRIM1 or control vector, with and without dox-induction, and treated with nerve growth factor for 5 days to induce neurite outgrowth (representative neurons shown in Figure 9d). The proportion of cells with neurites (defined as cellular process greater than cell body length)^66^ was quantified, as was neurite length. Expression of TRIM1 alone did not affect PC12 neurite outgrowth (49% +/- 2% neurite-bearing cells without TRIM1 expression; 46% +/- 9% neurite-bearing cells with TRIM1 expression, Figure 9e). LRRK2 G2019S induction significantly reduced the proportion of cells with neurite outgrowth, a phenotype that was completely rescued by co-expression of TRIM1 (29% +/- 6% neurite-bearing cells with LRRK2 G2019S without TRIM1; 44% +/- 6% neurite-bearing cells with LRRK2 G2019S and TRIM1). In cells bearing neurites, expression of TRIM1 without LRRK2 G2019S did not affect neurite length (40.8 um +/- 3.9 um without TRIM1 expression; 46.1 um +/- 8.5 um with TRIM1 expression, Figure 9f). LRRK2 G2019S induction reduced neurite length in the absence of TRIM1 while expression of TRIM1 with LRRK2 G2019S fully rescued neurite length (32.4 +/- 3.5 um with LRRK2 G2019S without TRIM1; 48.7 +/- 7.5 um with LRRK2 G2019S and TRIM1). Thus, in this model, TRIM1 protects against LRRK2 G2019S-induced neurite outgrowth defects.

## Discussion

Here we show that TRIM1 is a novel LRRK2 binding partner that ubiquitinates both WT and PD-mutant LRRK2 and influences LRRK2 degradation, subcellular localization, and Rab29 binding/kinase activation, as well as LRRK2 G2019S’s neurotoxic effects on neurite outgrowth (Figure 10). TRIM1 recruits LRRK2 to the microtubule cytoskeleton and drives LRRK2 K48-linked polyubiquitination and proteasomal degradation. Overexpression of TRIM1 decreased levels of overexpressed LRRK2 in cell lines and primary cortical neurons; knockdown and CRISPR knockout of endogenous TRIM1 increased steady-state levels of endogenous LRRK2. Until now, the ubiquitous protein CHIP was the only E3 ligase known to target LRRK2 to the proteasome.^20–22^ Interestingly, we found that TRIM1 drove LRRK2 proteasomal degradation more strongly than CHIP in our flow cytometric assay. CHIP mediates turnover of many unstable proteins,^22^ and appears particularly important for degradation of destabilized LRRK2 variants, such as the sporadic-PD risk allele, LRRK2 G2385R.^23^ One possible hypothesis is that CHIP may be especially important in degradation of unstable, misfolded LRRK2, while TRIM1’s role in LRRK2 degradation may be related to LRRK2’s phosphorylation state and possibly to LRRK2’s subcellular localization.

**Figure 10.**
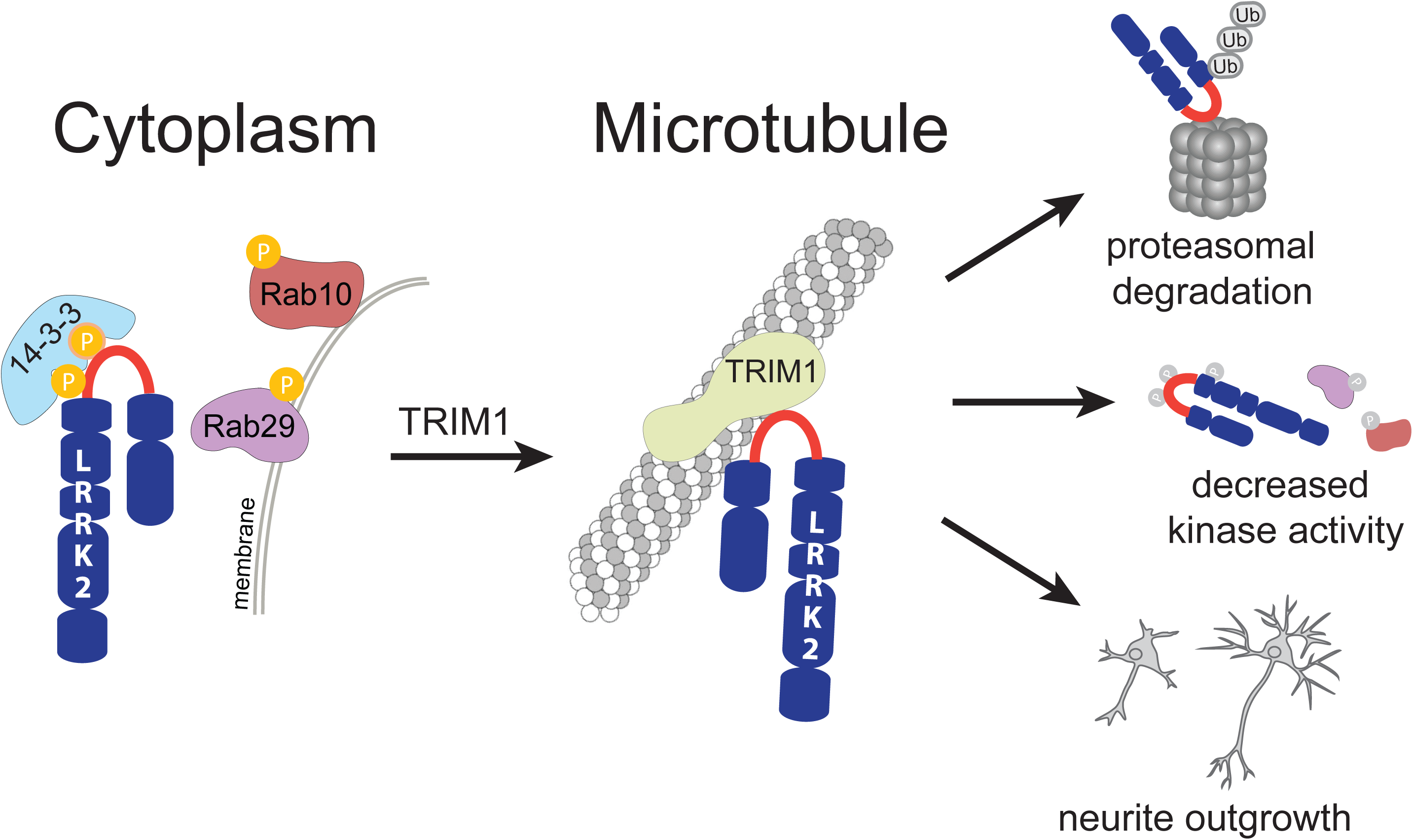
Model of the LRRK2 RL/TRIM1 interaction. Schematic highlighting the array of downstream effects of TRIM1 binding LRRK2 RL. Phosphorylation of S910 and S935 in the RL may act as a molecular switch, modulating LRRK2’s preference for binding 14-3-3 versus TRIM1, which in turn alters LRRK2 localization (cytoplasm versus microtubule), ubiquitination and turnover, kinase activity, and neuronal toxicity as measured by neurite outgrowth.

We used domain constructs to identify the regions mediating LRRK2’s interaction with TRIM1. TRIM1’s B-box domain (particularly B-box 1) was required for LRRK2 binding. LRRK2_822-982_, a 160 amino acid segment of LRRK2 between the ankyrin and LRR domains, was sufficient for TRIM1 to bind and microtubule-localize LRRK2. We utilized alanine scanning to further define the TRIM1 binding region of LRRK2, and found that only 9 amino acids, LRRK2_911-920_, are required for TRIM1 to bind, ubiquitinate, and bring LRRK2 to the microtubule. Interestingly, LRRK2_911-920_ lies between Ser910 and Ser935, both of which are phosphorylated by upstream kinases to allow LRRK2 to bind 14-3-3 proteins.^13, 33^ We find that the LRRK2-TRIM1 interaction can occur in the absence of Ser910/Ser935 phosphorylation, while, as others have demonstrated, the LRRK2-14-3-3 interaction requires phosphorylation of LRRK2 Ser910/Ser935. 14-3-3 binding stabilizes LRRK2 in the cytoplasm,^60^ whereas TRIM1 binding recruits overexpressed LRRK2 to microtubules. Interestingly, secondary structure modeling predicts LRRK2_822-982_ to be predominantly unstructured, consistent with the recent cryo-EM structure of full length LRRK2, for which LRRK2_853-981_ could not be solved.^31^ We therefore propose that LRRK2_853-981_ may serve as a regulatory loop (RL) whose phosphorylation status, and possibly conformation, may serve as a molecular switch dictating LRRK2’s binding to interacting partners. LRRK2’s close homolog LRRK1, which has a very similar domain structure to LRRK2, lacks the RL region. LRRK1 also contains no mutations linked to PD.^67^ We speculate that the RL region may be an important regulator of PD-relevant LRRK2 functions.

Like others, we find that Rab29 overexpression robustly augments WT and PD-mutant LRRK2 kinase function.^24^ Intriguingly, TRIM1 inhibits Rab29-mediated LRRK2 kinase activation (for both WT and PD-mutant R1441G LRRK2) and this effect is not due to decreased LRRK2 protein levels. Inhibition requires that TRIM1 contain an intact E3 ligase domain but does not require TRIM1 to bind microtubules, suggesting that ubiquitination but not microtubule-binding drives inhibition. We find that E3-intact TRIM1 augments the interaction between LRRK2 and Rab29, as measured by co-immunoprecipitation. This finding may suggest that ubiquitination of LRRK2 by TRIM1 changes the conformation of LRRK2 and causes LRRK2 to more strongly bind Rab29 in a kinase-inactive conformation. With 60% sequence coverage, we utilized MS to identify LRRK2 K831 as a site of TRIM1-mediated ubiquitination. However, a LRRK2 K831R mutation did not restore Rab29-mediated LRRK2 kinase activation in the presence of TRIM1. Because LRRK2 is such a large protein, 60% coverage identified 92 of 176 Lys residues but left the ubiquitination state of 84 Lys residues unknown. Thus, other ubiquitination sites on LRRK2 may be important for regulating the LRRK2-Rab29 interaction. Alternatively, we cannot rule out the possibility that TRIM1 ubiquitination of a non-LRRK2 target modulates the LRRK2-Rab29 interaction. TRIM1 ubiquitination sites on LRRK2 remain to be fully delineated as does the mechanism by which TRIM1 inhibits Rab29’s kinase activation of LRRK2.

This work uncovers the microtubule cytoskeleton as a potential site of LRRK2 turnover, a new role for the cytoskeleton in PD. Multiple groups have reported an association of LRRK2 with microtubules,^36–39^ including recent work that solved the structure of the LRRK2-microtubule interface to 14 Å,^37^ and a second recent study that suggests that LRRK2’s direct interaction with microtubules is regulated by the conformation of its kinase domain.^41^ However, the physiologic function of LRRK2’s microtubule association has not been rigorously investigated and some authors have suggested that LRRK2-microtubule filaments represent concentration-dependent “aggresomes.”^24^ We find that TRIM1, but not the highly homologous TRIM18, binds a 9 amino acid segment of LRRK2 to promote endogenous LRRK2 degradation. Although TRIM1 C, a microtubule non-binding construct, can ubiquitinate LRRK2, all published evidence suggests that endogenous TRIM1 and TRIM18 are entirely localized to microtubules. TRIM1 and TRIM18 are known to ubiquitinate other substrates at microtubules, targeting them for degradation. TRIM1 ubiquitinates astrin, a microtubule-associated protein involved in cell division, to decrease astrin levels and promote cell division.^68^ TRIM18 binds alpha4, a regulatory subunit of protein phosphatase 2a (PP2a), at microtubules, causing ubiquitination and degradation of PP2a.^69^ Our findings thus support a model in which microtubule association plays a physiologic role in LRRK2 biology.

Interestingly, while mCherry-TRIM1 coats microtubules in a uniform and smooth distribution (Figure 2b, left panel, inset), GFP-LRRK2 forms more discontinuous and punctate structures, possibly suggestive of multi-protein complexes (Figure 2b, middle panel, inset). We predict that, in addition to causing LRRK2 turnover, the association of TRIM1 and LRRK2 on the microtubule cytoskeleton is likely to have additional functions that are yet to be delineated. LRRK2’s phosphorylation of Rab8 and Rab10 plays a critical role in ciliogenesis^28, 70^ and centrosomal cohesion.^71^ Cilia and centrosomes are microtubule-based structures, and Rab8 and Rab10-positive vesicles intimately associate with them.^70^ One possibility is that endogenous LRRK2 undergoes regulated trafficking between physically proximate Rab-positive membranes, microtubules, and the cytoplasm. Because LRRK2 contains numerous protein-protein interaction domains and has many features of a scaffolding protein, identification of other LRRK2 binding partners at microtubules may provide important insight into additional functions and regulation, as well as potential therapeutic targets.

A limitation of our study is that we were not able to visualize endogenous LRRK2 via confocal microscopy and all microscopy studies utilized overexpressed LRRK2. In the cell types utilized in this work, the low levels of endogenously expressed LRRK2 have prohibited visualization of endogenous LRRK2 by confocal microscopy by any authors. In fact, the only study that we believe definitively identifies endogenous LRRK2 via confocal or other microscopy (Eguchi et al, 2018) utilizes macrophages, which express extremely low levels of TRIM1.^72, 73^ This limitation highlights an important roadblock in the field.^53^ Ongoing work by our group and others is focused on optimizing LRRK2 CRISPR tags, nanobodies, and other tools to allow definitive localization of LRRK2 and PD-mutant LRRK2.

Finally, TRIM1 falls within the PARK12 genomic locus on the X-chromosome.^46^ While it is tempting to speculate that TRIM1 mutations may be linked to PD, PARK12 is a large locus containing ∼600 genes and much additional work remains to determine if TRIM1 variants increase risk for PD. Regardless, TRIM1 may serve as a novel therapeutic target for PD, as suggested by our data that TRIM1 ameliorates LRRK2 G2019S-mediated neurite outgrowth defects. The mechanism by which LRRK2 G2019S inhibits neurite outgrowth is unknown; however, increases in both LRRK2 kinase activity and protein levels have been linked to neurotoxicity in PD,^35^ and we observe that TRIM1 regulates both. More broadly, our studies suggest that cellular pathways decreasing LRRK2 protein levels are possible targets to combat PD and should be identified and tested.

## Author Contributions

Conceptualization, G.S., N.J.K., S.A.O., A.H.; Methodology, A.E.D.S., F.S., M.F., E.M.E, H.A., E.V., D.L.S., G.S., J.V.H, T.P.M., J.R.J., J.V.D., C.M., T.W., R.J.N., T.C.C., S.F., E.J.D. A.H.; Investigation, , A.E.D.S., F.S., E.M.E., L.L., D.L.S, A.R., E.J.D., A.H.; Writing– Original Draft, A.H., S.A.O., A.E.D.S., A.H; Writing – Review & Editing, G.S., E.V., D.L.S, T.W., R.J.N., T.C.C., S.F., N.J.K.; Funding Acquisition, T.W., T.C.C., R.J.N., C.I., S.F., N.J.K., S.A.O., A.H.; Supervision, C.I., T.W., T.C.C., R.J.N., S.F., N.J.K., S.A.O., A.H.

## Acknowledgements

We thank Dario Alessi for use of his CRISPR edited GFP-LRRK2 A549 cell line, Daniela Boassa for her assistance with confocal microscopy, and David Coughlin and Michael Perry for assistance with statistical analyses. This work was made possible with support from NIH NINDS K08NS090633 (A.H.), DoD W81XWH-18-1-0376 (A.H.), the American Federation for Aging Research (A.H.), the PFCC and DRC center grant NIG P30 DK063720, NIH U54 HG008105 (S.F.), RF1AG058476 (S.F), Merck and Co. (S.F), the Head Start Program of the Michael J Fox Foundation (S.F.), the Taube/Koret Center (S.F), Michael J. Fox Foundation LRRK2 Challenge 2014 ID9550 (C.I.), Fondazione Banco di Sardegna grant number 2014.0489 (C.I.), Regione Sardegna grant number CRP-78083 (C.I), Fondo di Ateneo per la ricerca 2019 (C.I.), S10 RR026758 (T.W.), NIH R01EY027810 (S.A.O.), and NIH R01CA219815 (S.A.O.).

## CONTACT FOR REAGENT AND RESOURCE SHARING

Further information and requests for resources and reagents should be directed to and will be fulfilled by the Lead Contact, Annie Hiniker.

## MATERIALS AND METHODS

### Cell lines and tissue culture

All cell lines were grown at 37°C in a humidified atmosphere with 5% CO_2_. Human HEK-293T cells and human A549 cells were cultured in Dulbecco’s modified Eagle’s medium (DMEM) with 10% fetal bovine serum (FBS). Doxycycline-inducible GFP-LRRK2 HEK-293T cell lines^62^ were cultured in DMEM with 10% tetracycline-free FBS, 10 μg/ml blasticidin and 100 μg/ml hygromycin. Human H1299 cells were cultured in RPMI-1640 media with 10% FBS, 25 mM Hepes, and 2.0 g/L NaHCO_3_. Malme-3M cells were grown in RPMI-1640 with 10% FBS and 2 mM (1x) L-alanyl-L-glutamine (GlutaMAX, Thermo-Fisher). SK-N-SH human neuroblastoma cells were cultured in Eagle’s Minimum Essential Medium (EMEM). Rat primary cultures of cortical neurons were created from rat pup cortices at embryonic days 20-21 and cultured and differentiated as previously described^57^ in Neurobasal growth medium with 2mM GlutaMAX, Pen/Strep and B27 supplement (NB media). Rat PC-12 cells were grown in DMEM supplemented with 15% tetracycline-free FBS and 2mM GlutaMax. Doxycycline-inducible LRRK2 rat PC-12 cell lines^65^ were grown in DMEM supplemented with 10% horse serum and 5% tetracycline-free FBS or 15 % tetracycline-free FBS, 2 mM GlutaMAX, 100 U/ml penicillin G, 100 μg/ml streptomycin, 400 μg/ml G418, and 200 μg/ml hygromycin and were differentiated under low-serum conditions the same media containing 1% horse serum without FBS and 100 ng/ml NGF. PC12 cell differentiation media was replaced every 48 hours.

### Plasmids

Plasmids pcDNA5 frt/to expressing GFP-tagged human LRRK2, both full-length and truncation and point mutants and pCMV-C2-6myc or pCMV-C2-EGFP expressing WT human TRIM18 and TRIM1, both full-length and domain mutants have been previously described ^62, 74^ as has plasmid V5-14-3-3 theta ^27^. Plasmid expressing mCherry-tubulin was a gift from Roger Tsien ^75^, and plasmid pRK5-HA-ubiquitin WT was a gift from Ted Dawson (Addgene plasmid #17608)^76^.

Full-length human LRRK2 with N-terminal myc and FLAG tags was cloned into pcDNA5 frt/to. In brief, pCMV-2myc-LRRK2, a gift from Mark Cookson (Addgene plasmid #25361)^77^ was cloned into pcDNA5 frt/to by site-directed mutagenic removal of a single LRRK2 internal HpaI site (Quikchange, Stratagene), followed by HpaI/Eco53KI digest, ligation into EcoRV site, and return of HpaI site. A 2x FLAG tag was introduced upstream of the 2x myc tag by quikchange (Quikchange, Stratagene). Addgene plasmid #25361 was found to have the Arg50His variant not present in consensus Uniprot sequence (Q5S007) and site-directed mutagenesis was used to create Arg50.

Plasmid expressing GFP-LRRK2_822-982_ was created by introducing a stop codon in GFP-LRRK2_822-2527_ and plasmids expressing GFP-TRIM1_70-119_ (denoted “linker” Figure 4b), GFP-TRIM1_70-177_ (denoted “BB1”, Figure 4b), and GFPTRIM1_70-235_ (denoted “BB1,2”, Figure 4b) were created by introducing stop codons into GFP-TRIM1_ΔRF_. Western blot was used to verify that there was no read-through of the stop codon.

Plasmid expressing mCherry-myc-TRIM1 was created by cloning myc-TRIM1 into pmCherry-C1 (Clontech). Plasmid expressing EBFP2-14-3-3 theta was created using HiFi Cloning (NEB) of 14-3-3 theta into pEBFP2-C1 (Addgene plasmid #54665). A CHIP plasmid^78^ was a gift from Leonard Petrucelli and the CHIP ORF was cloned into pCMV-C2-6myc. Eos3.2-LRRK2 and mApple ORFs were synthesized and cloned into pGW1. All constructs were verified by DNA sequencing.

### Antibodies

The following antibodies were used: mouse anti-LRRK2 (Neuromab, N241A/34), rabbit anti-LRRK2 (Abcam, C41-2), rabbit anti-phospho-Ser910 LRRK2 (Abcam, UDD 1 15(3)), rabbit anti-phospho-Ser935 LRRK2 (Abcam, UDD2 10(12)), rabbit anti-phospho-Ser1292 LRRK2 (Abcam, ab203181), rabbit anti-FLAG (Sigma, F7425), mouse anti-myc (Sigma, clone 9E10, M4439), rabbit anti-myc (Cell Signaling, clone 71D10, 2278), rabbit anti-mCherry (Abcam, ab167453), rabbit anti-TRIM1 (Sigma, M2448; Thermo PA5-28457), mouse anti-GFP (ThermoFisher, clone GF28R, MA5-15256), rabbit anti-HA (Cell Signaling, clone C29F4, 3724S), mouse anti-HA (Sigma, H7411), mouse anti-V5 (ThermoFisher, clone E 10/V4RR, MA5-15253), mouse anti-Rab10 (Sigma, Sab5300028), rabbit anti-phospho-Thr71 Rab29 (Abcam, ab241062), rabbit anti-phospho-Thr73 Rab10 (Abcam, ab230261), mouse anti-ubiquitin (Millipore Sigma, MAB1510-I; Santa Cruz P4D1, sc-8017), rabbit linkage-specific K63 anti-ubiquitin (Abcam, ab179434), rabbit linkage-specific K48 anti-ubiquitin (Cell Signaling, 4289), rabbit anti-actin (Cell Signaling, clone 13E5, 4970), and mouse anti-actin (Sigma, A1978). For western blot, all primary antibodies were used at 1:250-1:1000 dilution except actin (1:1000-1:5000) and secondary antibodies (IRDye® 800CW or 680RD Goat anti-Mouse or anti-Rabbit IgG, LI-COR) were utilized at 1:10,000 dilution. For chemiluminescence of immunoblots, secondary antibodies used were peroxidase-conjugated AffiniPure Donkey anti-mouse (Jackson Immunoresearch 715-035-150) and peroxidase-conjugated Donkey anti-rabbit (Jackson Immunoresearch 711-035-152), with SuperSignal West Femto Maximum Sensitivity Substrate (Thermo, 34094). For immunoprecipitation and co-immunoprecipitation, the following pre-conjugated agarose-resin systems were used according to manufacturer instructions: anti-FLAG M2 (Sigma, A2220); GFP-trap®_A or magnetic GFP-trap®_MA, myc-trap®_A or magnetic myc-trap®_MA or mCherry-trap®_A (Chromotek); Pierce anti-HA (Thermo), magnetic anti-V5 beads (MBL International), and magnetic-bead conjugated panselective TUBE2 (LifeSensors, UM402M). Anti-TRIM1 antibodies were conjugated to magnetic dynabeads following manufacturer’s instructions (Invitrogen).

### Cell transfection, drug treatment, and lysis

HEK-293T cells were transfected using Fugene 6 (Promega), Lipofectamine LTX, or Lipofectamine 2000 (Thermo-Fisher); H1299 cells were transfected with Lipofectamine 2000; and PC12 cells were transfected with Lipofectamine LTX or Lipofectamine 3000, all per manufacturer’s instructions. LRRK2 expression was induced with 2 ng/ml to 1 μg/ml doxycycline. The proteasomal inhibitor MG132 was used at 2 μM for 24 hours. Autophagy was inhibited with chloroquine at 25 μM for 24 hours. LRRK2 kinase inihibitor MLi-2 was used at 100-200 nM for 24 hours for protein turnover assays and 4 hours for colocalization studies, and at 500 nM for 5 hours for co-immunoprecipitation studies. Proteasome inhibitor bortezomib was used at 1-100nM for 8-18 hours. For cell lysates, cells were harvested by scraping or trypsinization, pelleted and washed with PBS 2x, and lysed either by pipetting up and down or by homogenization using a pellet pestle (Kimble Scientific) in 1 ml ice-cold immunoprecipitation buffer (50 mM Tris pH 7.5, 150 mM NaCl, 1 mM EDTA, and 0.5% NP-40) supplemented with cOmplete protease inhibitor (Roche, 11836170001) and Phosstop phosphatase inhibitor (Roche, 04906845001) and rotated end over end at 4°C for 30 minutes and debris pelleted at 10,000-15,000 rcf for 5 min at 4°C. Lysate total protein concentration was measured using the Pierce BCA Protein Assay Kit (Thermo Scientific, 23225).

For rat cortical neurons, at 3–4 days *in vitro*, neurons were transfected with plasmids and Lipofectamine 3000. Prior to adding the lipofectamine-DNA mix, cells were incubated in Neurobasal with 1x KY media (10 mM Kynurenic acid, 0.0025% phenol red, 5 mM HEPES and 100 mM MgCl_2_). Primary neurons were incubated in lipofectamine-DNA mix for 20–40 minutes, washed with Neurobasal, and cultured in NB medium.

### Immunoprecipitation, mass spectrometry, and data analysis of LRRK2 interactome

6 µg of either pcDNA5 containing the above FLAG-LRRK2 construct or FLAG tag alone was transfected into HEK-293T cells using Fugene 6. Cells were harvested 48 hours after transfection using 10 mM EDTA in D-PBS for 5 minutes at 25°C, followed by 2 PBS washes. Cells were lysed as above, with protein concentrations normalized by BCA assay, and the resulting lysates incubated with 30 µl anti-FLAG M2 (Sigma, A2220) pre-conjugated agarose-resin for 2-3 hours at 4°C. FLAG affinity purification followed by mass spectrometry was carried out essentially as described in Jager et al. except that elution of bound proteins was done using 100 μg/ml FLAG peptide (Sigma, F3290).^44^ In brief, following elution, 10 µl of the IP eluate were reduced with 2.5 mM DTT at 60°C for 30 minutes followed by alkylation with 2.5 mM iodoacetamide for 40 minutes at room temperature. 100 ng sequencing grade modified trypsin (Promega) was then added to the sample and incubated overnight at 37°C. Peptides were then desalted on ZipTip C18 pipette tips (Millipore), lyophilized to dryness, and resuspended in a final solution of 0.1% formic acid for injection into a Thermo Scientific LTQ Orbitrap XL Mass Spectrometer. For AP-MS experiments, raw data conversion and Protein Prospector search were also performed as described previously.^44^

### Genome editing of TRIM1

CRISPR-Cas9 mediated knock-out of the TRIM1 genomic locus in HEK-293T cells was performed essentially as described.^79^ A single guide RNA (sgRNA) targeting the N-terminus of TRIM1’s RING domain (5’-CAACTCTAGGCAGATTGGAC-3’) with low off-targeting score was chosen following careful transcript analysis using NCBI and the Zhang lab CRISPR Design tool (crispr.mit.edu). Complementary dsDNA oligos with BbsI-compatible overhangs were designed, dsDNA guide inserts ligated into BbsI-digested pSpCas9(BB)-2A-GFP (PX458) (AddGene plasmid #48138), and plasmid verified by sequencing. 40 hours prior to transfection, HEK-293T cells were seeded at 4 x 10^5^ cells per 35 mm well. Transfection of 2.5 µg PX458/sgRNA was performed using Lipofectamine 3000. 24-48 hours post-transfection, cells were harvested and GFP-positive cells were isolated using a BD FACSAria II and grown in DMEM supplemented with 10% FBS. After two weeks, GFP-negative cells were single-cell sorted (BD FACSAria II) into 96-well plates and single clones were expanded. After reaching ∼80% confluency, individual clones were transferred into six-well plates and then expanded further and genomic DNA harvested as below.

For genetic verification of knockout, genomic DNA was isolated using a KAPA Mouse Genotyping Kit (Roche). PCR amplification of the N-terminal region of TRIM1 was performed using Q5 High-Fidelity 2X Master Mix (New England BioLabs) with the primers: F=5’-TGTGTTCAGCACAGAAATGCCT-3’, R=5’-AGGCAGGCTTAGAATTAG CCC-3’), cloned into the TOPO TA cloning kit (Thermo), and single colonies sequenced. For each cloning reaction, DNA was isolated from >8 bacterial colonies using QIAprep® Spin Miniprep Kit (Qiagen) and TRIM1 knockout confirmed by sequencing using a T3 forward promoter to confirm truncating stop codons in all copies of the TRIM1 gene present in the genome of selected clones. Isolation of genomic DNA and sequencing from the parental WT HEK-293T cell line was performed in parallel and demonstrated wild-type alleles of TRIM1 as expected. Absence of endogenous TRIM1 protein was confirmed using immunoprecipitation with anti-TRIM1 antibodies conjugated to dynabeads followed by immunoblotting for TRIM1 in parallel to the positive-control parental wild type cells (Figure 1e; see immunoprecipitation protocol below).

### Immunoprecipitation and co-immunoprecipitation (co-IP)

Cells were lysed in immunoprecipitation buffer as above and lysate protein concentrations quantified by BCA and normalized prior to addition of antibody-conjugated beads 20-40 µl antibody-conjugated beads (GFP, myc, HA, or V5, listed above, or dynabead-conjugated anti-TRIM1) were added and immunoprecipitations were performed at 4°C for 2-12 h. Beads were washed ≥3x in wash buffer (50 mM Tris pH 7.5, 150-500 mM NaCl, 1 mM EDTA) and bound proteins eluted by heating at 70-95°C in 40-80 µl 4x NuPage LDS loading buffer with 5% β-mercaptoethanol. Samples were run on NuPage Novex mini gels, either 4-12% Bis-Tris or 3-8% Tris-Acetate, then transferred onto PVDF membrane with the Genscript eBlot L1 transfer system and blocked with LI-COR Intercept TBS blocking buffer. All imaging and quantification of immunoblots was performed using a LI-COR Odyssey® CLx imaging system and Image Studio Lite software version 5.2. All quantification of LRRK2 or phospho-LRRK2 protein levels was performed using 3-8% Tris-Acetate gels.

### Quantification of LRRK2 protein levels in TRIM1 CRISPR KO

WT HEK-293T and TRIM1 Knockout CRISPR HEK-293T cells were seeded at 3×10^6^ cells per plate on 10cm plates. After 24 hours, cells were harvested and lysed in immunoprecipitation buffer (50 mM Tris pH 7.5, 150 mM NaCl, 1 mM EDTA, and 0.5% NP-40) supplemented with cOmplete protease inhibitor (Roche) and Phosstop phosphatase inhibitor (Roche). Cells were lysed by pipetting and cell lysates were rotated at 4°C for 30 minutes then debris pelleted at 4°C at 5,000 rpm for 10 min. Protein levels in the lysates were normalized, then samples were run on a Tris-Acetate gel at 100 volts for 110 minutes.

### Isolation of Endogenously Ubiquitinated LRRK2 using Tandem Ubiquitin Binding Entities

HEK-293T cells were seeded in 10 cm dishes and transfected with FLAG-LRRK2 in the presence of myc-TRIM1 or myc alone vector control. After transfection for 8 hours, cells were treated with 100 nM bortezomib for 12 hours. 20 hours post-transfection, cells were lysed as above and lysate protein concentrations normalized using BCA assay. FLAG-LRRK2 immunoprecipitation was performed using 30 μl FLAG antibody-conjugated beads (Sigma) at 4°C for 1 h. FLAG-LRRK2 was eluted by incubation with 100ug/mL 3x FLAG peptide at 4°C for 1 h. 50 μl TUBE2 magnetic beads (LifeSensors) was washed 3x in TBS-T, added to eluted FLAG-LRRK2, and rotated at 4°C for overnight. TUBE2 beads were washed 3x (50 mM Tris pH 7.5, 250 mM NaCl, 0.2 % NP-40, 1 mM DTT) and bound proteins eluted by heating at 70200°C for 10 minutes in 20 μl 4x Nupage LDS loading buffer (Invitrogen, NP0007) containing 5% β-mercaptoethanol. Standards used were K48 and K63 linked recombinant polyubiquitin chains (R&D Biosystems).

### LRRK2 siRNA knockdown and co-immunoprecipitation with endogenous TRIM1

800 nM of either LRRK2 siRNA (Dharmacon Cat #M-006323-02-0010) or control scrambled siRNA (#D-0001206-13-05) was electroporated into HEK-293T cells according to the Lonza Kit V protocol. After 48 hours growth at 37°C, cells were harvested and lysed as above for co-IP. 50 µl antibody-conjugated dynabeads (cat #14311D), previously coupled to TRIM1 antibody (Thermo, PA5-28457) or IgG isotype control (BD Pharmingen, 550326), were incubated with lysates for 16 hours at 4°C. Samples were washed, eluted, and immunoblotted as above.

### Live cell confocal imaging

Cells were seeded onto poly-D-lysine–coated glass-bottom 35 mm dishes (MatTek Corporation), with media and transfection conditions as described above. Spinning disk and laser scanning confocal live cell imaging was performed under environmentally controlled conditions, at 37°C and 5% CO_2_. For experiments utilizing the spinning disk microscope, the protocol is as described in Stehbens et al^80^ except that the system was upgraded with a next generation scientific CCD camera (cMyo, 293 Photometrics) with 4.5 um pixels allowing optimal spatial sampling using a 60x NA 1.49 objective (CFI 294 APO TIRF; Nikon). For experiments using the Olympus Fluoview 1000 laser scanning confocal microscope, a 60X oil immersion objective with numerical aperture 1.42 was utilized to obtain confocal images (1024 x 1024 pixels). Z-stack images were acquired with a step size of 0.5-1 μm and processed using the Fiji software package.

### Quantification of microtubule-bound LRRK2

H1299 cells were seeded at a density of 4 x10^5^ cells per plate on 35mm glass-bottom dishes (MatTek) and transfected with indicated plasmids using Lipofectamine 2000. 24 hours after transfection, live cell imaging was performed under environmentally controlled conditions, at 37°C and 5% CO_2_. Cells were imaged at 60x using either a Nikon Eclipse Ti Fluorescence microscope or a Olympus Fluoview 1000 laser scanning confocal microscope, using a 60X oil immersion objective. Cells were only selected for analysis if they fluoresced in both the red and green channels and expressed TRIM1 or tubulin at microtubules. Each cell was categorized based on whether LRRK2 localized to microtubules or not with reviewer blinded to construct transfected. 50 cells were counted per plate with two plates per transfection condition, for a total of 100 cells per experiment. This was repeated for a total of three independent experiments. For each condition, the number of cells per replicate with LRRK2 localized at microtubules were averaged and expressed as a percentage with error was calculated as the standard deviation.

### Flow Cytometry

GFP-LRRK2 levels were measured in doxycycline-inducible cell lines on a Fortessa flow cytometer (BD Biosciences). Cells were induced with 2-5 ng/ml doxycycline, and doxycycline was washed from cells 18-72 hours prior to analysis. On the day of analysis, cells were trypsinized and pelleted and washed before being resuspended in DPBS. GFP intensity was measured using a 488 nm laser for excitation, and a detector with 505-nm long pass filter and a 525/50-nm band pass filter. Only live, single cells, as determined by forward and side scatter, were analyzed.

### CRISPRi knockdown of TRIM1

Catalytically dead Cas9 (dCas9-BFP) was inserted randomly into doxycycline-inducible GFP-LRRK2 HEK-293T cells via lentiviral transduction. Cells were sorted for a BFP+ pure population on an Aria2SORP. BFP intensity was measured using a 405 nm laser for excitation, and a detector with a 450/50-nm band pass filter. The top four predicted guide RNAs for TRIM1 based on *Horlbeck et al.*^81^ were packaged with lentiviral vectors and added to cells, and then puromycin-selected (0.75 μg/mL) for 2 days before cells were plated and induced for included experiments. Knockdown was measured via real-time quantitative PCR with a non-targeting guide RNA used as a control.

### siRNA knockdown of TRIM1

800 nM of either TRIM1 siRNA (Dharmacon Cat #L-006938-00-0005) or control scrambled siRNA (#D-0001206-13-05) was electroporated into Malme-3M cells according to the Lonza Kit V protocol. After 24 hours incubation, cells were harvested and RNA was extracted following the NucleoSpin RNA Plus kit protocol (Macherey-Nagel) for qPCR. A TaqMan Gene Expression Assay probe against TRIM1/MID2 (Life Technologies Corporation) was used to confirm the knockdown of TRIM1 mRNA. After 48 hours, a parallel replicate of each knockdown was harvested, lysed, and immunoblotted for LRRK2 to measure the effect of TRIM1 knockdowns on LRRK2 protein levels. Protein levels were visualized using two antibodies against LRRK2, C41-2 and N241.

### Identification of ubiquitinated LRRK2 lysines by MS

HEK293T cells were seeded in a 15 cm cell culture dish and transfected with FLAG-LRRK2 and HA-ubiquitin plasmids in the presence of myc-TRIM1, myc-ΔRF TRIM1, or myc alone vector control. Sequential immunoprecipitation for FLAG and HA were performed on lysates as described above. Protein samples were subsequently reduced and alkylated, and digested with trypsin overnight at 37°C. Peptides were then desalted on C18 ziptip columns, lyophilized to dryness, and resuspended in 0.1% formic acid for injection into an Orbitrap Fusion Lumos Tribrid Mass Spectrometer. Raw data was analyzed with MaxQuant to identify and quantify LRRK2 ubiquitination at K831, as well as ubiquitin chain abundance.^82^ Quantification across samples was normalized by LRRK2 protein abundance.

### LRRK2 turnover assay with robotic microscope imaging system

Primary rat cortical neurons in 96-well plates were co-transfected with pGW1-GFP and pGW1-mEos3.2-LRRK2, and either TRIM1 or control plasmid. To measure the degradation of LRRK2 neurons expressing pGW1-GFP, pGW1-Eos3.2-LRRK2, and either TRIM1 or empty vector were photoswitched with a 5–8 s pulse of light at 405-nm wavelength 30-36 hours after transfection. Prior to photoswitching neurons transfected with mEos3.2-LRRK2 only fluoresce green with no detectable red fluorescence; however, upon photoswitching, a population of the green protein is irreversibly switched to emit red fluorescence. We then imaged the cells for red fluorescence (mEos3.2-LRRK2-red) every 4-10 hours for the next 2 days. Custom-based automated algorithms were used to capture images of neurons in each group in a high-throughput and unbiased manner. Live transfected neurons were selected for analysis based on pGW1-GFP fluorescence intensity and morphology. Neurons were only selected if they had extended processes at the start of the experiment. The abrupt loss of pGW1-GFP fluorescence was used to estimate the survival time of the neuron ^57, 83^. The Eos3.2-LRRK2-red fluorescence intensity was measured in each individual neuron at each time point that it was alive using a region of interest that corresponded to the cell soma. The co-transfected pGW1-GFP provided the morphology mask for the cell soma.

### Measurement of phosphorylation in the presence of Rab29 and TRIM1

Doxycycline-inducible GFP-LRRK2 HEK-293T cell lines were cultured as above on 60 mm dishes and transfected at 70% confluence with 0.5 µg pAH116 (myc-TRIM1) and/or 0.5 µg pAH358 (HA-Rab29) for 6 hrs using Lipofectamine 2000. After 6 hrs, the transfection media was removed and replaced with media containing 1 μg/mL doxycycline. Cells were incubated in dox-containing media for 18 hours, and harvested and lysed as above. Lysate total protein concentrations were measured by BCA and 30 μg of lysate was loaded per sample. Gel electrophoresis, transfer, immunoblotting, and quantification was performed as described above.

### PC12 neurite outgrowth assay

Doxycycline-inducible LRRK2 G2019S PC12 cells were plated at 20,000 cells/well in 96-well plates and transfected with pmaxGFP at 30 ng/well (for analysis of cell morphology) and mCherry-TRIM1 or mCherry empty vector at 200 ng/well. 24 hours after transfection, cells were moved to poly-d-lysine coated cover slips (Neuvitro) in the presence and absence of 1 µg/ml doxycycline. 48 hours after transfection, media was changed to PC12 differentiation media +/- 1 µg/ml doxycycline. Cells were then grown in differentiation media for 5 days with media changed every 48-60 hours. Cells were then fixed in 4% paraformaldehyde-PBS for 20 min, washed 3x with PBS, permeabilized in PBS with 10% goat serum, 0.4% Triton X-100, 30 mg/ml BSA and 10 mg/ml glycine for 1 hour, washed 3x with PBS, and mounted on slides using Vectashield hardmount with DAPI. All steps from fixation to mounting were performed at 25°C. Cells were imaged at 40X using a Keyence BZ-X700 fluorescence microscope; all cells containing both red and green fluorescence were imaged. The presence/absence of neurites and length of neurites was assessed using ImageJ. Dead cells were excluded from further analysis.

## QUANTIFICATION AND STATISTICAL ANALYSIS

### Statistical analysis

General statistical analysis was performed using excel, R, python, or STATA. For analysis of flow cytometry data, significance was evaluated using either an unpaired, two-tailed Student’s t-test for two-sample comparisons or ANOVA with post-hoc t-test and Bonferroni correction for three or more groups. For data for which normality could not be assumed, non-parametric testing was used, either the Mann-Whitney U test for two-sample comparison or Kruskal-Wallis followed by post-hoc Dunn test with Bonferroni correction for multiple comparisons. To test the effect of TRIM1 on the presence or absence of neurite outgrowth, we modelled the proportion of cells with neurites using a binomial distribution (which approximates a normal distribution at our sample size) and tested the null hypothesis that the groups had the same probability of having neurites using a Z-test with Bonferroni correction. A p-value < 0.05 was considered statistically significant.

Proteins were identified using Protein Prospector, and high-confidence protein-interactions were identified by label-free quantification of bait samples as compared to empty vector control using MSstats R-package ^45^. Two independent experiments with two or three independent replicates of WT FLAG-LRRK2 compared to FLAG empty vector were included in the analysis. In the case of rare proteins in which peptides were seen in the presence of LRRK2 but none were identified in the empty vector control, a Mann-Whitney U test was performed to identify proteins significantly increased in the LRRK2 sample.

Flow cytometry data was analyzed using FlowJo software (FlowJo LLC). Data represent the normalized median green fluorescence intensity and twice the standard error of the mean.

To measure the degradation of LRRK2 in the robotic microscope imaging system, Red-LRRK2 fluorescence was measured longitudinally in each cell for at least 48h or until its death. The Red-LRRK2 intensity values from each cell were fitted to an exponential and used to derive a LRRK2 half-life value. Cells were excluded from the analysis if the Red-LRRK2 intensity values were lower than local background intensity. Cells without a monotonic decrease in Red-LRRK2 signal or with a half-life greater than three standard deviations from the mean were also excluded. The majority of these excluded cells were due to out of focus images. ANOVA analysis was used to compare significant differences between mean half-lives across the groups.

### Data Availability

The authors declare that the data supporting the findings of this study are available within the paper [and its supplementary information files].

---

**Table S1, related to Figure 1. LRRK2 interacting partners.**

**Table S2, related to Figure 3. LRRK2 peptides and ubiquitination sites identified.**

**Movie S1, related to** **Figure 2. Time-lapse of GFP-LRRK2 localization in the presence of mCherry-TRIM1.**

## Supplemental Figure Titles and Legends

**Figure S1.**
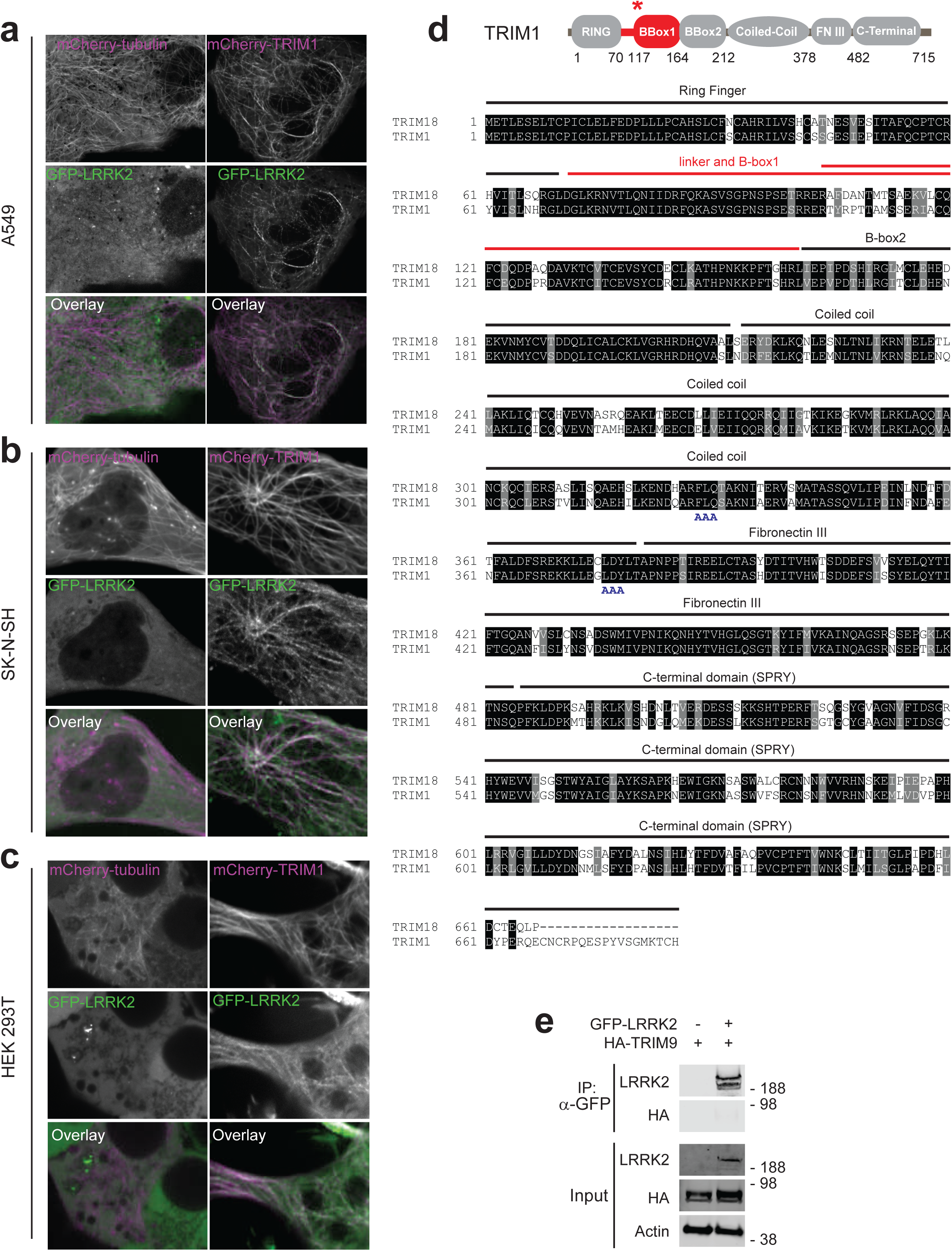
Additional characterization of the LRRK2-TRIM1 interaction. Live-cell confocal microscopy of GFP-LRRK2 and mCherry-tubulin or mCherry-TRIM1 transiently transfected into (A) A549 cells, (B) SK-N-SH cells, or (C) HEK-293T cells. From top to bottom, each set shows mCherry-tubulin or mCherry-TRIM1, GFP-LRRK2, merged image. In all lines examined, in the presence of mCherry-tubulin, GFP-LRRK2 is diffusely cytoplasmic, but microtubule localized in the presence of mCherry-TRIM1. (D) Alignment of TRIM18 with TRIM1. Domains labeled above alignment. Red line designates region required for TRIM1 interaction with LRRK2. Double red line designates region of least homology in TRIM1 and TRIM18 dual B-box domain. Dual-AAA motifs below the sequence designate the mutated amino acids used to make cytoplasmic TRIM1 C variant. (E) Immunoprecipitation of GFP-LRRK2, which fails to co-immunoprecipitate with HA-TRIM9 in HEK-293T cells.

**Figure S2.**
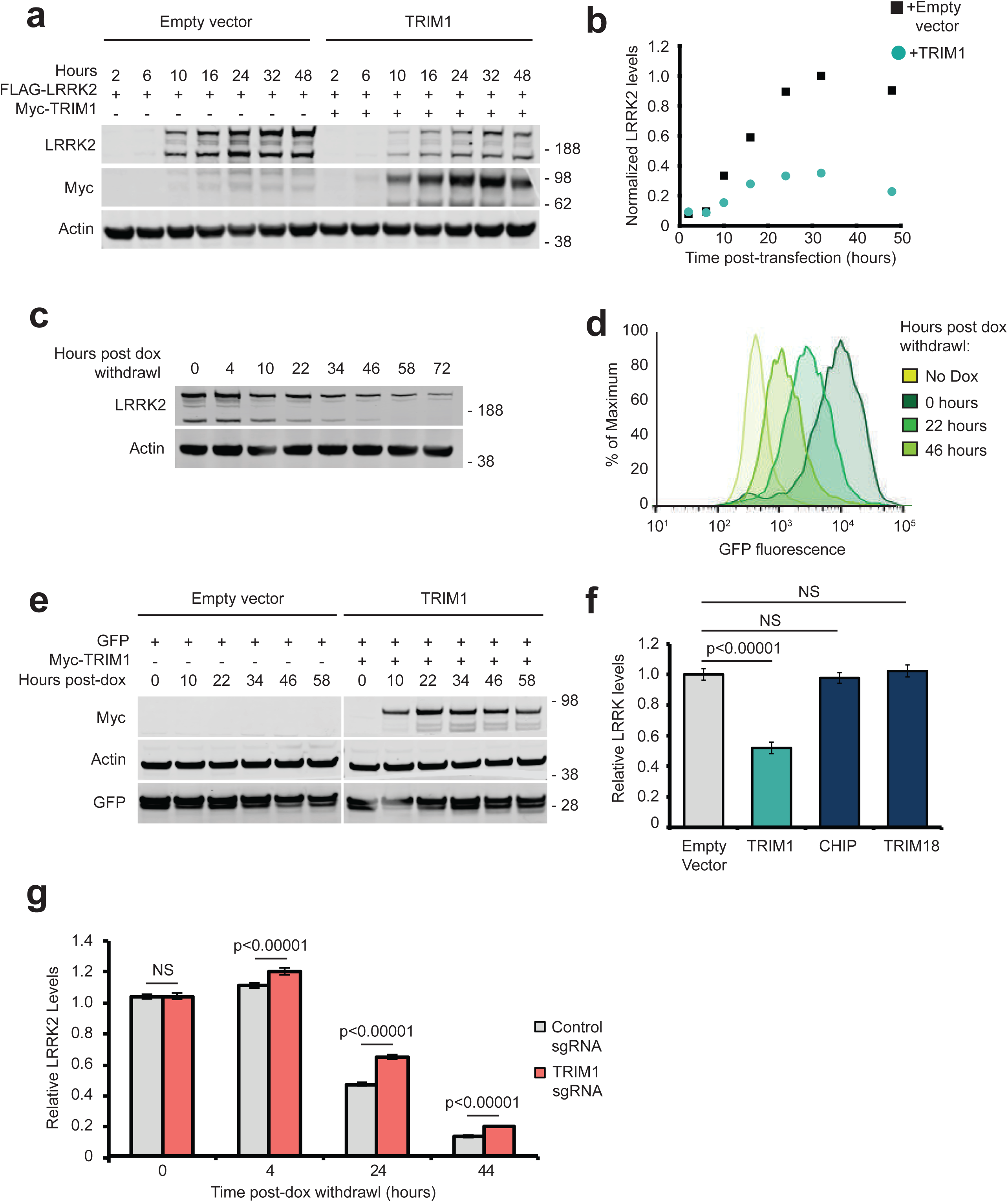
Evaluation of LRRK2 levels and validation of flow cytometric system to measure LRRK2 turnover. (A) Immunoblot of FLAG-LRRK2 co-transfected with myc-TRIM1 or empty vector control. Time indicates hours after transfection. (B) Quantification of panel (A) with LRRK2 levels normalized to actin. (C) Immunoblot showing LRRK2 levels relative to actin after withdrawal of doxycycline (doxycycline-induced for 18 hours). (D) Histograms of GFP fluorescence from samples immunoblotted in (C). (E) Immunoblot of dox-induced GFP expression co-transfected with myc-TRIM1 or empty vector control. Time indicates hours after transfection. (F) Flow cytometric quantification of GFP-LRRK2 levels in the presence of TRIM1, CHIP, or TRIM18. (G) Flow cytometric quantification of GFP-LRRK2 levels in TRIM1 knockdown and control dCas9/dox-GFP-LRRK2 HEK-293T lines at 0, 4, 24, and 44 hours after dox withdrawal relative to 0 hours. Significance testing for panels F and G was performed using ANOVA with post-hoc t-test with Bonferroni correction.

**Figure S3.**
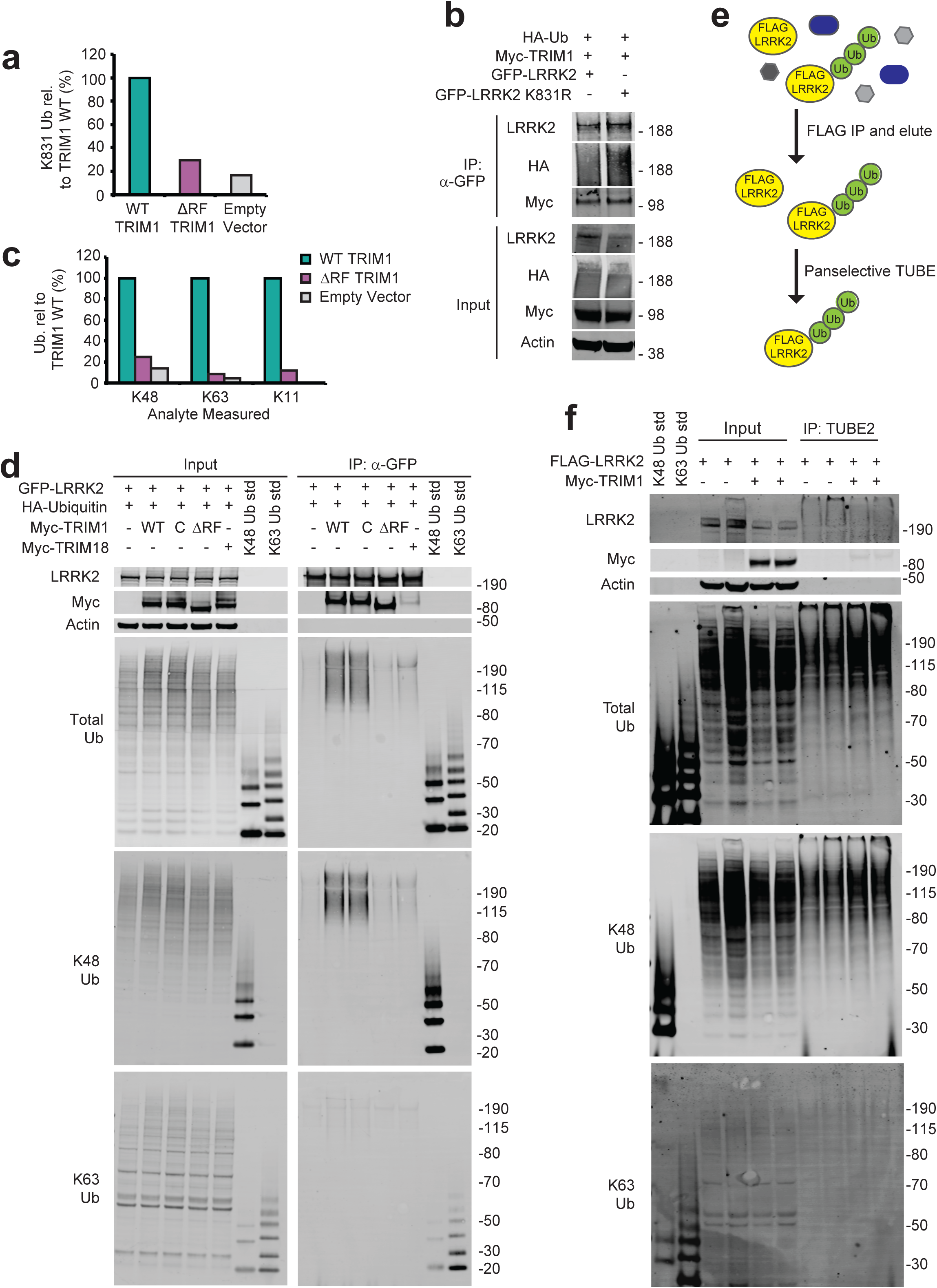
LRRK2 ubiquitination by TRIM1. (A) Quantitative MS analysis of LRRK2 K831 ubiquitination in the presence of WT TRIM1, ΔRF TRIM1, or empty vector. (B) Co- immunoprecipitation and ubiquitination of GFP-LRRK2 WT or K831R with myc-TRIM1 in the presence of HA-ubiquitin in HEK-293T cells. (C) All ubiquitin linkages identified by MS analysis of ubiquitinated LRRK2 eluate in the presence of WT TRIM1, ΔRF TRIM1, or empty vector. (D) Co-immunoprecipitation and ubiquitination of GFP-LRRK2 in the presence of HA-Ubiquitin and TRIM1 WT, C, or ΔRF or TRIM18 followed by immunoblotting against total ubiquitin, K48-linked ubiquitin, or K63-linked ubiquitin. (E) Schematic of tandem ubiquitin binding entity (TUBE) assay. LRRK2 was immunoprecipitated from HEK-293T lysate with anti-FLAG-conjugated agarose resin, and then a panselective TUBE was used to isolate ubiquitinated LRRK2, which was analyzed by immunoblot. (F) TUBE assay with FLAG-LRRK2 expressed in the presence of myc-TRIM1 or a control vector. TUBE eluates were blotted with broad anti-ubiquitin antibodies as well as K48 and K63 linkage-specific antibodies. All immunoblots are representative of at least three independent experiments.

**Figure S4.**
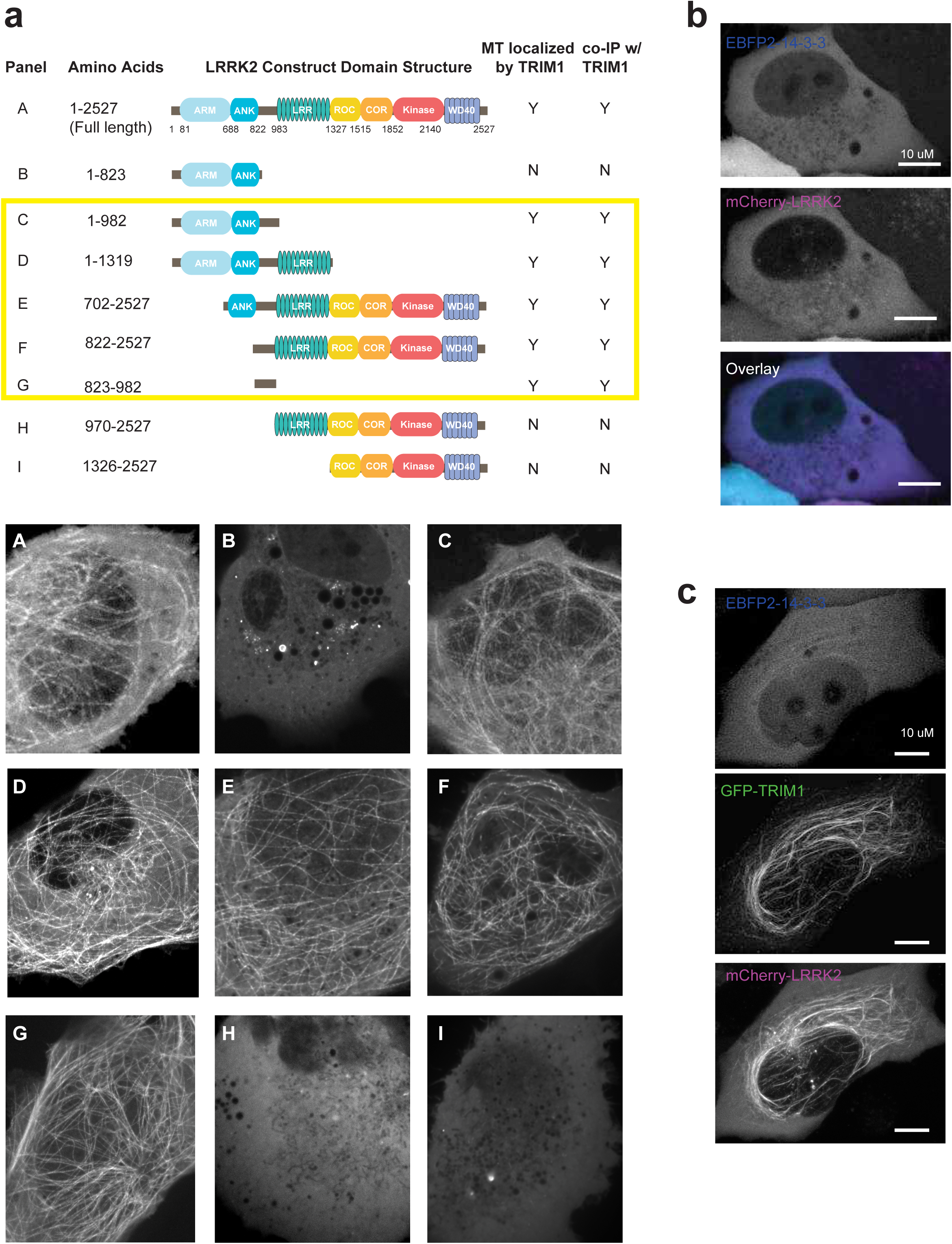
TRIM1 binds LRRK2-RL to cause LRRK2 microtubule localization. (A) Schematic of GFP-LRRK2 constructs (above) with corresponding live-cell microscopy in the presence of mCherry-TRIM1 in H1299 cells (below). Each panel shows only fluorescence at 488 (GFP) to illustrate the subcellular localization of each GFP-LRRK2 construct in the presence of mCherry-TRIM1, which is always localized to the microtubule network. (B) Live-cell confocal microscopy of mCherry-LRRK2 in the presence of EBFP2-14-3-3 in H1299 cells. (C) Live-cell confocal microscopy of mCherry-LRRK2 in the presence of EBFP2-14-3-3 and GFP-TRIM1 showing individual EBFP2, GFP, and mCherry channels from Figure 6A.

**Figure S5.**
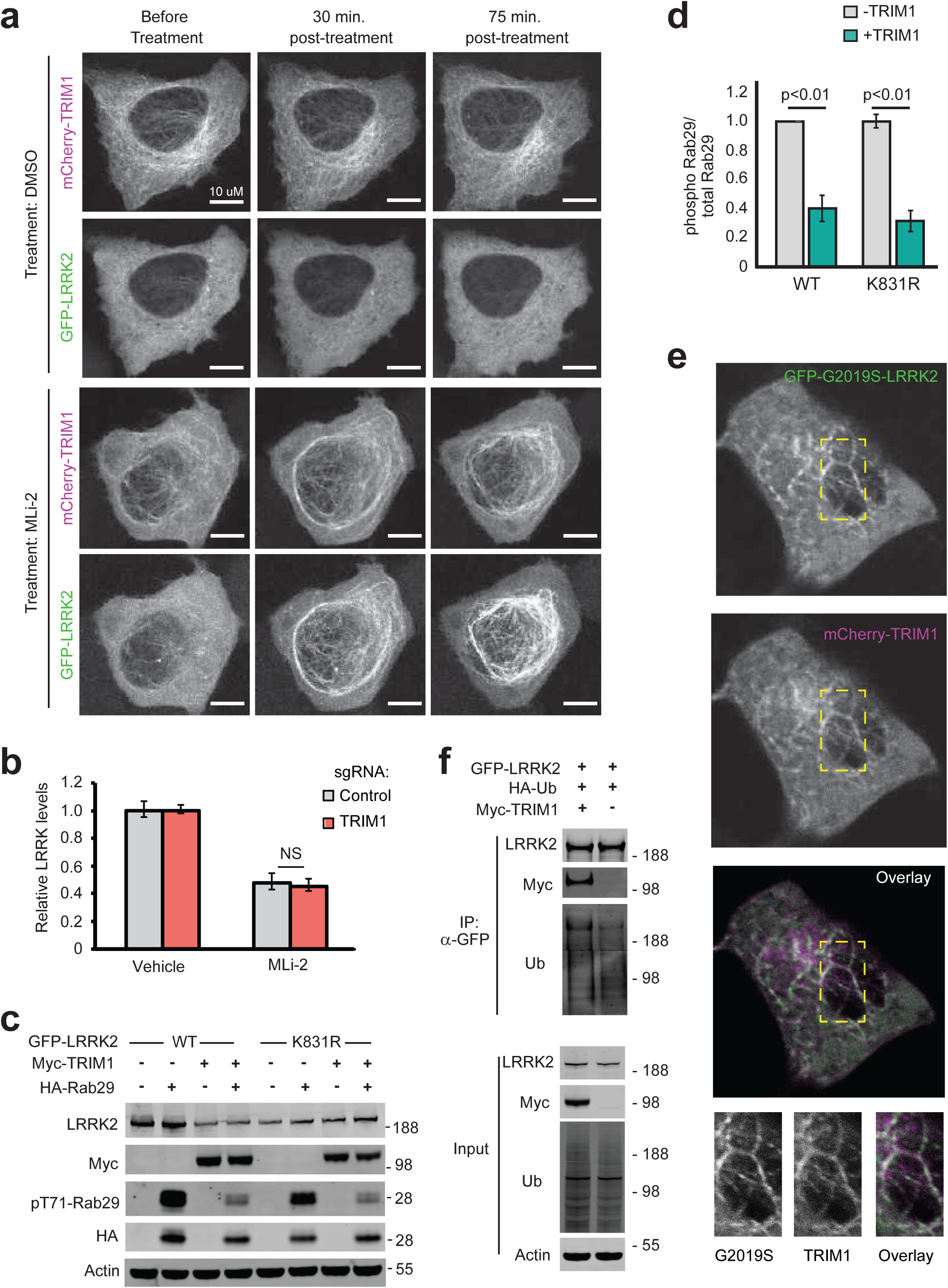
Characterization of LRRK2 co-expressed with 14-3-3 or cytoplasmic TRIM1. (A) Live-cell confocal microscopy of GFP-LRRK2 in the presence of mCherry-TRIM1 following treatment with LRRK2 kinase inhibitor MLi-2 (200 nM) or vehicle, showing the individual mCherry and GFP channels from time course in Figure 6G. Rare cells with low levels of co-localization prior to treatment were followed over time. (B) Quantification of GFP-LRRK2 fluorescence in flow cytometric assay with TRIM1 knocked down (red bar) compared to cells with non-targeting sgRNA (grey bar). Cells were dox-induced for 24 hours, dox was removed and MLi-2 (100 nM) or vehicle was added for another 24 hours before cells were assayed. (C) Immunoblot of Rab29 phosphorylation in the presence and absence of TRIM1 for wild type LRRK2 and LRRK2 K831R. (D) Quantification of Rab29 phosphorylation in (C). (E) Live-cell confocal microscopy of GFP-LRRK2 G2019S and mCherry-TRIM1 transiently transfected into PC-12 cells. From top to bottom: GFP-LRRK2 G2019S, mCherry-TRIM1, merged image, higher magnification of area in yellow boxes. (F) Immunoprecipitation and ubiquitination of GFP-LRRK2 with myc-TRIM1 in the presence of HA-ubiquitin in PC12 cells. Significance testing for panel B was performed using ANOVA with post-hoc t-test with Bonferroni correction and for panel D was performed using Mann-Whitney U test.

